# A neuropsychiatric disease-associated mutation in LRRC8B disrupts cellular Ca²⁺ signaling, mitochondrial function, and bioenergetics

**DOI:** 10.64898/2026.04.16.718892

**Authors:** Athira Ajith, Durai Shalu, Rudrakant Sharma, Amit Kumar Ghosh, Amal Kanti Bera

**Affiliations:** Department of Biotechnology, Bhupat and Jyoti Mehta School of Biosciences, Indian Institute of Technology Madras, Chennai - 600036, Tamil Nadu, India

**Keywords:** LRRC8B, ER calcium leak, mitochondrial calcium uptake, mitochondrial dysfunction, SMI

## Abstract

Leucine-rich repeat-containing 8 (LRRC8) proteins form the volume-regulated anion channel (VRAC) and participate in diverse physiological processes, including cell volume regulation, gliotransmitter release, and insulin secretion. In mammals, five paralogs (LRRC8A–E) exist; LRRC8A is the obligatory subunit that assembles into functional hexameric channels with LRRC8C, D, or E. LRRC8B is distinct: we previously demonstrated its role in regulating endoplasmic reticulum (ER) Ca²⁺ homeostasis and ER Ca²⁺ leak. A LRRC8B variant (Y380S) identified in an Indian family with severe mental illness has been associated with disease pathology, but its molecular and cellular consequences remain unknown. Here, we show that this disease-associated mutant perturbs Ca²⁺ signalling, mitochondrial bioenergetics, and redox homeostasis. Both wild-type and mutant LRRC8B localize to the ER and mitochondria. LRRC8B knockdown significantly reduced mitochondrial Ca²⁺ uptake and maximal respiratory the Y380S mutant phenocopied LRRC8B knockdown, altering ER Ca²⁺ release, elevating basal cytosolic Ca²⁺, and impairing mitochondrial Ca²⁺ uptake, consistent with a dominant-negative mechanism. The mutant further induced mitochondrial dysfunction, including loss of membrane potential, oxidative stress, and defective antioxidant responses, ultimately compromising cellular bioenergetics and viability. Mechanistically, the Y380S mutation disrupted LRRC8B interaction with the mitochondrial outer membrane channel VDAC. These findings identify LRRC8B–VDAC coupling as a key determinant of mitochondrial Ca²⁺ handling and provide a mechanistic link between LRRC8B dysfunction and neuropsychiatric disease.

**Highlights:**

- A psychiatric disease–associated LRRC8B variant (Y380S) acts as a dominant-negative regulator of ER Ca²⁺ homeostasis. It enlarges the releasable ER Ca²⁺ pool and reduces cell viability.
- LRRC8B promotes mitochondrial Ca²⁺ uptake through interaction with VDAC. The Y380S mutation disrupts this interaction, reducing mitochondrial Ca²⁺ uptake.
- The Y380S mutant increases mitochondrial superoxide production without activating compensatory antioxidant responses.
- The mutant also causes mitochondrial membrane depolarization and bioenergetic failure, as evidenced by reduced oxygen consumption rate and ATP production.

**Graphical abstract:** 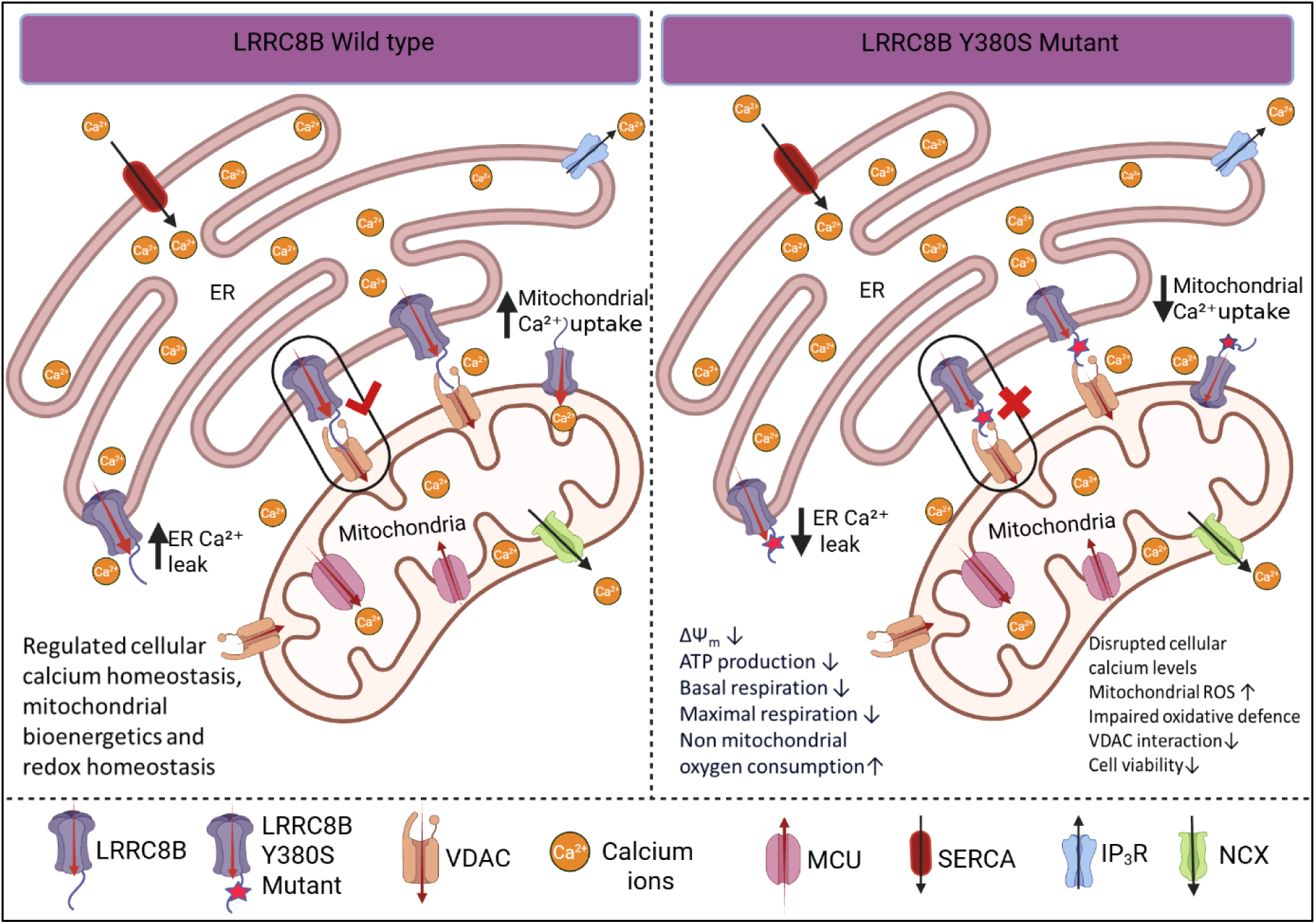

## Introduction

The leucine-rich repeat containing 8 (LRRC8) gene was first identified in a patient with congenital agammaglobulinemia carrying a balanced translocation t(9;20)(q33.2;q12), which disrupted a previously uncharacterized gene on chromosome 9 (Miyake et al., 1995; Sawada et al., 2003). Subsequent studies identified this locus as LRRC8A, which encodes a protein with four transmembrane domains and a C-terminal leucine-rich repeat (LRR) region (Kubota et al., 2004). The translocation results in truncation of LRRC8A, deleting the eighth, ninth, and part of the seventh LRR domains near the C-terminus, and generates a chimeric transcript co-expressed with the intact protein from the unaffected allele. Four additional paralogs, LRRC8B, LRRC8C, LRRC8D, and LRRC8E, were later identified, forming the LRRC8 protein family within the broader LRR superfamily (Abascal & Zardoya, 2012; Kubota et al., 2004).

In 2014, two independent studies established LRRC8 proteins as the molecular components of the volume-regulated anion channel (VRAC), a ubiquitously expressed chloride channel activated by cell swelling to mediate regulatory volume decrease (Qiu et al., 2014; Voss et al., 2014). Functional VRACs are heteromeric complexes composed of the essential subunit LRRC8A together with one or more paralogous subunits such as LRRC8C, LRRC8D, or LRRC8E (Karakas et al., 2025; Yanushkevich et al., 2025). The specific subunit composition determines key channel properties, including gating kinetics, regulatory mechanisms, and substrate selectivity (Syeda et al., 2016). Notably, unlike other paralogs, there is no convincing evidence that LRRC8B forms functional VRAC channels. In primary astrocytes, siRNA-mediated knockdown of LRRC8B did not affect VRAC-mediated efflux of aspartate and taurine under hypotonic conditions, whereas knockdown of other subunits markedly reduced osmolyte efflux (Schober et al., 2017). Similarly, reconstitution studies in *Xenopus* oocytes showed that LRRC8A forms functional channels with LRRC8C, LRRC8D, and LRRC8E, whereas the ability of LRRC8B to form functional channels was not established (Gaitán-Peñas et al., 2016). Consistent with this, engineered HEK293T cells lacking all LRRC8 paralogues except LRRC8A and LRRC8B fail to exhibit measurable VRAC currents, whereas LRRC8A in combination with LRRC8C, LRRC8D, or LRRC8E forms functional channels (Lutter et al., 2017). Together, these findings indicate that LRRC8B is dispensable for VRAC activity and likely serves a distinct cellular function.

Indeed, LRRC8B differs from other family members in both its subcellular localization and function. It localizes predominantly to the endoplasmic reticulum (ER) and mitochondria (Ghosh et al., 2017). Our previous work demonstrated that LRRC8B mediates passive Ca²⁺ efflux from the ER, thereby regulating luminal Ca²⁺ levels and modulating store-operated calcium entry (SOCE). Specifically, LRRC8B overexpression reduces ER Ca²⁺ content and enhances SOCE, whereas its silencing delays ER Ca²⁺ depletion following SERCA inhibition (Ghosh et al., 2017). These findings establish LRRC8B as an important regulator of ER Ca²⁺ leak and intracellular Ca²⁺ homeostasis.

Efficient Ca²⁺ transfer from the ER to mitochondria is critical for mitochondrial bioenergetics. This process occurs at ER–mitochondria contact sites, where the voltage-dependent anion channel (VDAC) on the mitochondrial outer membrane cooperates with the inositol 1,4,5-trisphosphate receptor (IP3R) on the ER membrane (Marchi et al., 2014; Rizzuto et al., 2012; Szabadkai et al., 2006). VDAC serves as the primary entry route for Ca²⁺ into mitochondria and regulates mitochondrial Ca²⁺ uptake through interactions with ER proteins (De Stefani et al., 2012). Given its localization to both the ER and mitochondria, LRRC8B is well positioned to modulate this Ca²⁺ transfer pathway.

LRRC8B is highly expressed in the brain (Pervaiz et al., 2019), where dysregulation of Ca²⁺ signaling can impair bioenergetics, increase oxidative stress, and promote neuroinflammation—processes strongly linked to neurological disorders (Mustaly-Kalimi et al., 2025; Zündorf & Reiser, 2011). Whole-exome sequencing of patients from an Indian family identified a rare LRRC8B variant (Y380S) associated with severe mental illness (SMI) (Ganesh et al., 2019). However, it remains unclear whether this variant alters or antagonizes wild-type LRRC8B function and what downstream cellular consequences it may have.

In this study, we demonstrate that LRRC8B not only regulates ER Ca²⁺ leak but also physically interacts with VDAC to control mitochondrial Ca²⁺ uptake, thereby acting as a key mediator of ER–mitochondrial Ca²⁺ crosstalk. The Y380S mutation disrupts ER Ca²⁺ release and mitochondrial Ca²⁺ uptake, leading to abnormal Ca²⁺ distribution, loss of mitochondrial membrane potential, increased oxidative stress, impaired antioxidant responses, defective respiration, and reduced cell viability. Furthermore, we show that the Y380S mutation weakens the interaction between LRRC8B and VDAC, providing a mechanistic basis for impaired mitochondrial Ca²⁺ handling. Together, these findings reveal a broader role for LRRC8B in cellular Ca²⁺ homeostasis and highlight its potential relevance in SMI.

## Materials and Methods

### Plasmids

The plasmid encoding GFP-tagged human LRRC8B (HG24935-ACG; Sino Biological Inc, USA). The LRRC8B-Y380S mutant was generated from this construct via site-directed mutagenesis and sequence-verified. The pEGFP-N1 vector (6085-1; Clontech, USA) served as the fluorescent empty-vector control. Subcellular Ca²⁺ dynamics were assessed using compartment-specific genetically encoded calcium indicators obtained from Addgene, USA. ER Ca²⁺ levels were monitored using the ER-targeted red fluorescent Ca²⁺ indicator R-CEPIA1er (58216), originally deposited by Dr Masamitsu Iino (Suzuki et al., 2014). Mitochondrial Ca^2+^ was measured using R-CEPIA3mt (140464), also deposited by Dr Masamitsu Iino (Kanemaru et al., 2020), and cytosolic Ca^2+^ was assessed with the genetically encoded indicator cyto-RCaMP1h (105014) gifted by Dr Franck Polleux (Hirabayashi et al., 2017).

### Antibodies and chemicals

Primary antibodies used in this study included anti-LRRC8B (HPA017950; Sigma-Aldrich, USA) and anti-β-actin (A5441; Sigma-Aldrich, USA), along with anti-VDAC (D73D12; Cell Signalling Technology, USA) and anti-GFP (SC9996; Santa Cruz Biotechnology, USA). HRP-conjugated anti-rabbit IgG (7074), HRP-conjugated anti-mouse IgG (7076), and Normal Rabbit IgG (2729) were obtained from Cell Signalling Technology, USA

The fluorescent dyes Fura-2-AM (50033), ER-Tracker Red (87917), Mito Tracker Red (M7512), and Mitosox (M36008) were purchased from Invitrogen Life Technologies (USA). Hoechst 33342 (B2261), tetramethylrhodamine ethyl ester (TMRE) (87917), Histamine (H7250), thapsigargin (T9033), ionomycin (I0634), 3-(4,5-dimethylthiazol-2-yl)-2,5-diphenyl tetrazolium bromide (MTT) (475989), phenylmethylsulfonyl fluoride phenylmethylsulfonyl fluoride (PMSF) (P7626) and protease inhibitor cocktail (P8340) were sourced from Sigma-Aldrich (USA). All other chemicals and reagents were of analytical grade and obtained from Sigma-Aldrich (USA) unless otherwise stated.

### Cell culture and transfection

HEK293T cells were obtained from the National Centre for Cell Science (Pune, India) and cultured in Dulbecco’s Modified Eagle Medium (DMEM) supplemented with 10% heat-inactivated fetal bovine serum (FBS) and 1X antibiotic–antimycotic solution (15240-062; Gibco, USA) at 37°C in a humidified atmosphere containing 5% CO₂. For imaging experiments, cells were seeded onto glass coverslips in 35 mm dishes, while 60 mm dishes were used for biochemical assays. All transfections were carried out at 50–60% confluency using JetPrime transfection reagent (01000046; Polyplus, USA) with 2–3 µg of total plasmid DNA for a 35 mm dish. In co-expression and overexpression experiments, an empty pEGFP-N1 vector was included to normalize total DNA content across all experimental groups. For LRRC8B silencing, cells were transfected with a validated siRNA duplex targeting human LRRC8B (5′-GUCAUCUUAUCACAUCUUA-3′; SR-NP001-020, Eurogentec, Belgium) at a final concentration of 25 nM. A scrambled non-targeting siRNA (Mission siRNA Universal Negative Control; cat. no. SIC001, Sigma-Aldrich, USA) was used at 100 nM as the negative control.

### Mitochondrial fractionation

Mitochondria were isolated following the protocol of (Huang et al., 2012) with minor modifications. Briefly, HEK293T cells were collected by centrifugation at 100 × g for 5 minutes, washed with ice-cold phosphate-buffered saline (PBS), and re-suspended in ice-cold mitochondrial isolation buffer (225 mM mannitol, 75 mM sucrose, 0.1 mM EGTA, 30 mM Tris-HCl, pH 7.4) containing 0.1 mg/ml PMSF and a protease inhibitor cocktail.

Cells were gently homogenized and subjected to low-speed centrifugation at 1,300 × g for 5 minutes at 4°C to remove cell debris. The resulting supernatant was then centrifuged at 12,000 × g for 25 minutes at 4°C to sediment the mitochondrial fraction. The pellet was subsequently re-suspended in freshly prepared RIPA buffer supplemented with PMSF and protease inhibitor cocktail, followed by vortexing for 5 minutes to ensure complete lysis. Enrichment of the mitochondrial fraction was verified by western blot analysis using an antibody targeting VDAC as a mitochondrial marker.

### Immunoblotting

To assess endogenous expression, knockdown efficiency, and overexpression of LRRC8B, protein levels were analyzed by Western blotting. Whole-cell lysates or organellar fractions were prepared by resuspension in RIPA buffer (10 mM Tris–HCl (pH 7.4),1 mM EDTA, 1% TritonX-100, 0.1% SDS, 0.1% Sodium Deoxycholate,140 mM NaCl) supplemented with PMSF and a protease inhibitor cocktail, followed by brief vortexing. Protein concentrations were determined using the Bradford assay. Equal amounts of protein were denatured in 6× Laemmli sample buffer and resolved on 10 % SDS-polyacrylamide gels. Proteins were subsequently transferred onto PVDF membranes, which were blocked with 5% BSA and incubated with primary antibodies overnight at 4°C (1:1000 dilution). Membranes were then washed and incubated with HRP-conjugated secondary antibodies (1:5000 dilution) for 1 h at room temperature. Immunoreactive bands were detected using Clarity Western ECL substrate (1705060; Bio-Rad, USA) and visualized using Image Lab software (Bio-Rad). β-actin was used as a loading control in all Western blot experiments.

### Confocal microscopy

HEK293T cells were seeded in glass-bottom dishes and transiently transfected with plasmids encoding GFP-tagged wild-type (WT) or mutant LRRC8B. At 24 h post-transfection, cells were incubated with 1 µM ER-Tracker Red or MitoTracker Red for 20–30 min at 37°C to label the ER or mitochondria, respectively. Cells were subsequently washed with 1× PBS and counterstained with 2 µg/ml Hoechst 33342 for 10 min at room temperature to visualize nuclei. Following a final wash, cells were imaged using an Olympus FluoView 3000 laser-scanning confocal microscope.

### Electrophysiological Recording of VRAC Currents

VRAC currents were recorded using the whole-cell configuration of the patch-clamp technique. Patch pipettes were fabricated from borosilicate glass capillaries (Sutter Instruments, USA) using a P-97 horizontal puller and had resistances of 2 to 5 MΩ when filled with pipette solution. Pipette capacitance and series resistance were compensated prior to recording. The isotonic extracellular (bath) solution contained (in mM): 140 NaCl, 6 CsCl, 2 CaCl₂, 2 MgCl₂, 6 glucose, and 10 HEPES, adjusted to pH 7.4 with NaOH. The intracellular (pipette) solution contained (in mM): 140 CsCl, 1 MgCl₂, 5 EGTA, 4 Na₂ATP, and 10 HEPES, adjusted to pH 7.2 with CsOH. CsCl was used as the primary intracellular salt to suppress endogenous K⁺ currents and improve voltage-clamp quality. EGTA was included to chelate residual Ca²⁺ and prevent Ca²⁺-activated Cl⁻ channel activation. After establishing the whole-cell configuration, baseline currents were recorded under isotonic conditions using a voltage-step protocol (holding potential of −60 mV; steps from −100 to +100 mV in 10 mV increments; 500 ms per step). VRAC currents were subsequently activated by perfusion of a hypotonic bath solution, prepared by reducing NaCl from 140 to 90 mM while keeping all other constituents unchanged. After 5 min of continuous hypotonic challenge, a duration sufficient for maximal VRAC activation, currents were re-recorded from the same cell using an identical voltage-step protocol. All recordings were performed using an Axopatch 200B amplifier (Molecular Devices, USA). Analog signals were low-pass filtered at 2 kHz and digitized at 10 kHz. Data acquisition and analysis were performed using pCLAMP 10 software and Clampfit 10 (Molecular Devices, USA), respectively. VRAC currents were isolated by subtraction of the isotonic (baseline) current traces from those recorded under hypotonic conditions at each corresponding voltage step. Current densities (current amplitude/ cell capacitance) were derived from the net VRAC current amplitudes and plotted as a function of membrane voltage.

### Cell viability assay

Cell viability was assessed using the MTT assay. Briefly, cells were incubated with MTT for the 2 hours, allowing metabolically active cells to reduce the yellow tetrazolium salt to insoluble purple formazan crystals via mitochondrial dehydrogenase activity. Following incubation, the medium was removed and formazan crystals were solubilized in dimethyl sulfoxide (DMSO). Absorbance was measured at 590 nm using a microplate reader, with background subtraction at an appropriate reference wavelength. Cell viability was calculated as a percentage relative to vector-transfected control cells, which were set to 100%. All measurements were performed in technical triplicate and repeated in at least three independent experiments.

### Ca²⁺ measurements

For imaging experiments, HEK293T cells were seeded onto 12 mm coverslips. Organelle-specific Ca²⁺ sensor plasmids were co-transfected with either LRRC8B-WT or mutant overexpression plasmids, or with siRNA targeting LRRC8B. Cells were briefly washed in HBSS (140 mM NaCl, 5.6 mM KCl, 3.6 mM NaHCO₃, 1.8 mM CaCl₂, 1 mM MgCl₂, 5.6 mM glucose, 10 mM HEPES, pH 7.4) and imaged using an Olympus IX83 fluorescence microscope with appropriate filter sets. Histamine (100 µM) was used to evoke ER Ca²⁺ release, 1 μM thapsigargin (TG) was used to inhibit ER Ca²⁺ refilling, and 5 μM ionomycin was used to elevate cytosolic Ca²⁺ levels in the respective experiments. ER Ca²⁺ levels were measured using RCEPIA1er, while cytosolic Ca²⁺ was monitored using either the genetically encoded sensor CytoRCaMP or the ratiometric dye Fura-2 AM. Mitochondrial Ca²⁺ dynamics were recorded using the mitochondria-targeted sensor RCEPIA2mt.

### Mitochondrial potential and ROS measurement

Mitochondrial potential (ΔΨ_m_) was assessed using TMRE. Cells were incubated with TMRE (200 nM) for 5 min at room temperature in the dark, followed by two washes with HBSS. Fluorescence images were acquired using an Olympus IX83 microscope. For real-time depolarization measurements, FCCP (1 µM), an uncoupler of mitochondrial oxidative phosphorylation, was applied during imaging. Mitochondrial ROS was measured with MitoSOX™ Red, which predominantly detects superoxide radicals. HEK293T cells were incubated in 5 µM MitoSOX™ Red in serum-free media and placed in the dark for 20 minutes at 37°C in a CO_2_ incubator. Cells were washed with 1X PBS twice and imaged to evaluate mitochondrial ROS with microscopy using RFP filter.

### mRNA expression analysis

Transfected HEK293T cells were washed with ice-cold PBS, and total RNA was extracted using the TRIzol reagent (9108; Takara, Japan). Briefly, cells were lysed in TRIzol, scraped, and transferred to 1.5 ml microcentrifuge tubes. Phase separation was achieved by adding chloroform followed by centrifugation. The aqueous phase was collected, and RNA was precipitated with isopropanol, washed with 70% ethanol, and dissolved in RNase-free water. First-strand cDNA synthesis was performed using the Bio-Rad iScript cDNA Synthesis Kit (170-8891; Bio-Rad, USA) according to the manufacturer’s instructions. Quantitative real-time PCR (qPCR) was carried out using gene-specific primers (Table S1) at a final concentration of 200 nM. Relative gene expression was calculated using the 2^−ΔΔCt method, with β-actin serving as the endogenous reference gene.

### Seahorse XF assay

Mitochondrial function was assessed in live HEK293T cells using the Seahorse XF96 Analyzer (Agilent Technologies). Cells (5 × 10⁴ per well) were seeded in XF96 cell culture microplates under the indicated conditions. Prior to measurement, cells were washed twice with Seahorse assay medium (DMEM supplemented with 200 mM glutamine, 100 mM pyruvate, and 1 M glucose) and equilibrated in 180 µl of pre-warmed medium at 37°C in a non-CO₂ incubator for 45 min. Mitochondrial respiration was evaluated using the XF Cell Mito Stress Test Kit (103015-100; Agilent Technologies, USA) according to the manufacturer’s protocol. Oligomycin (1.5 µM), FCCP (1 µM), and rotenone/antimycin A (0.5 µM each) were sequentially injected to assess ATP-linked respiration, maximal respiratory capacity, and non-mitochondrial oxygen consumption, respectively. The oxygen consumption rate (OCR) was normalized to total protein content per well, determined by BCA assay, to enable comparison across experimental groups.

### Co-immunoprecipitation

For co-immunoprecipitation, confluent HEK293 cells were washed twice with ice-cold PBS, scraped, and lysed in 1× RIPA buffer supplemented with 0.1 mg/mL PMSF and protease inhibitor cocktail. Cell lysates were transferred to pre-chilled 1.5 ml tubes, gently agitated for 45 minutes at 4 °C, and centrifuged at 13,000 × g (Hermle, Germany) for 25 minutes at 4 °C. The supernatants were collected into fresh pre-chilled tubes, and protein concentration was determined using Bradford reagent (Bio-Rad Laboratories, USA). For each reaction, 1 mg of total protein was incubated with anti-GFP at a 1:500 dilution and rotated overnight at 4 °C. Protein A/G PLUS-agarose (30 µL; sc-2003, Santa Cruz Biotechnology, USA) was washed with ice-cold PBS and added to the antibody–lysate mixture, followed by rotation for an additional 3 hours at 4 °C. The beads were then washed three times with ice-cold PBS and re-suspended in 15 µl of Lämmli sample buffer. Samples were heated at 95 °C for 10 minutes and briefly centrifuged. Proteins were resolved by SDS–PAGE and analyzed by immunoblotting as indicated. All co-immunoprecipitation experiments were performed in triplicate. As a negative control, lysates containing 1 mg of total protein were incubated with an isotype-matched Normal Rabbit IgG (2729; CST, USA) (same concentration as GFP antibody) followed by Protein A/G PLUS-agarose under identical conditions. IgG control immunoprecipitates were processed in parallel and analyzed by SDS–PAGE and immunoblotting.

### LC–MS/MS analysis of pulled-down samples

After co-immunoprecipitation, beads were washed with 50 mM Tris-HCl (pH 7.5) and processed for proteomic analysis. Proteins were subjected to on-bead digestion in 2 M urea, followed by reduction with DTT, alkylation with iodoacetamide, and overnight trypsin digestion as described above. Peptides were acidified to approximately 1% (v/v) formic acid, desalted using C18 StageTips, eluted with 70% acetonitrile in 0.1% formic acid, dried in a vacuum concentrator, and reconstituted in water containing 0.1% formic acid for LC–MS/MS analysis.

Peptides were analyzed by liquid chromatography–tandem mass spectrometry (LC–MS/MS) on an Orbitrap Exploris 200 mass spectrometer (Thermo Fisher Scientific, USA) coupled to a nanoLC system under standard proteomics conditions. Data were acquired in data-dependent acquisition (DDA) mode, and label-free quantification (LFQ) was performed at the MS1 level. Raw LC–MS/MS data were processed using FragPipe v23.1, with database searching performed using MSFragger against the UniProt *Homo sapiens* proteome database. Peptide and protein identifications were filtered to a false discovery rate of 1% using Philosopher, and label-free quantification was performed using IonQuant. Differential protein abundance and visualisation of volcano plots were performed using FragPipe Analyst.

### Image analysis and data extraction

For imaging experiments, regions of interests (ROIs) corresponding to individual cells were manually selected in CellSens software following background subtraction. Fluorescence intensity for each ROI was extracted across time frames and exported to Microsoft Excel for analysis. In time-lapse experiments, baseline fluorescence (F_₀_) was defined as the average intensity immediately before agonist application. Changes in fluorescence (F) were normalized to baseline, and responses were expressed as fold change (F/F_₀_) over time. The peak of each trace was taken as the maximum fold change. In selected analyses, half-time (t_₁/₂_) values were determined as the time required to reach 50% of the maximal (or minimal) fold change from baseline.

### Statistical analysis

All statistical analyses were performed using GraphPad Prism 11. Data are presented as mean ± SEM. Differences between two groups were assessed using an unpaired two-tailed Student’s t-test for normally distributed data, and comparisons across more than two groups were evaluated by one-way ANOVA followed by Tukey’s post-hoc test. A P-value of less than 0.05 was considered statistically significant. Significance levels are denoted as follows: *p < 0.05, **p < 0.01, ***p < 0.001, and ****p < 0.0001. N denotes independent biological replicates and n represents individual cells

## Results

### LRRC8B-Y380S retains normal subcellular localization but reduces cell viability

In our previous work, we demonstrated that LRRC8B localizes primarily to the ER and mitochondria. To determine whether the Y380S mutation affects this distribution, we expressed GFP-tagged wild-type LRRC8B or LRRC8B-Y380S in HEK293T cells and examined subcellular localization by confocal microscopy. The ER, mitochondria, and nuclei were labelled with ER-Tracker red, MitoTracker red, and Hoechst, respectively. Both wild-type and mutant LRRC8B showed substantial co-localization with each organelle marker, as evidenced by the yellow signal in merged images (Fig 1A). Quantitative analysis using CellSens software on background-subtracted images yielded mean Pearson’s correlation coefficients of 0.709 for the ER and 0.604 for mitochondria. These results demonstrate that the Y380S mutation does not alter the subcellular localization of LRRC8B.

**Figure 1.**
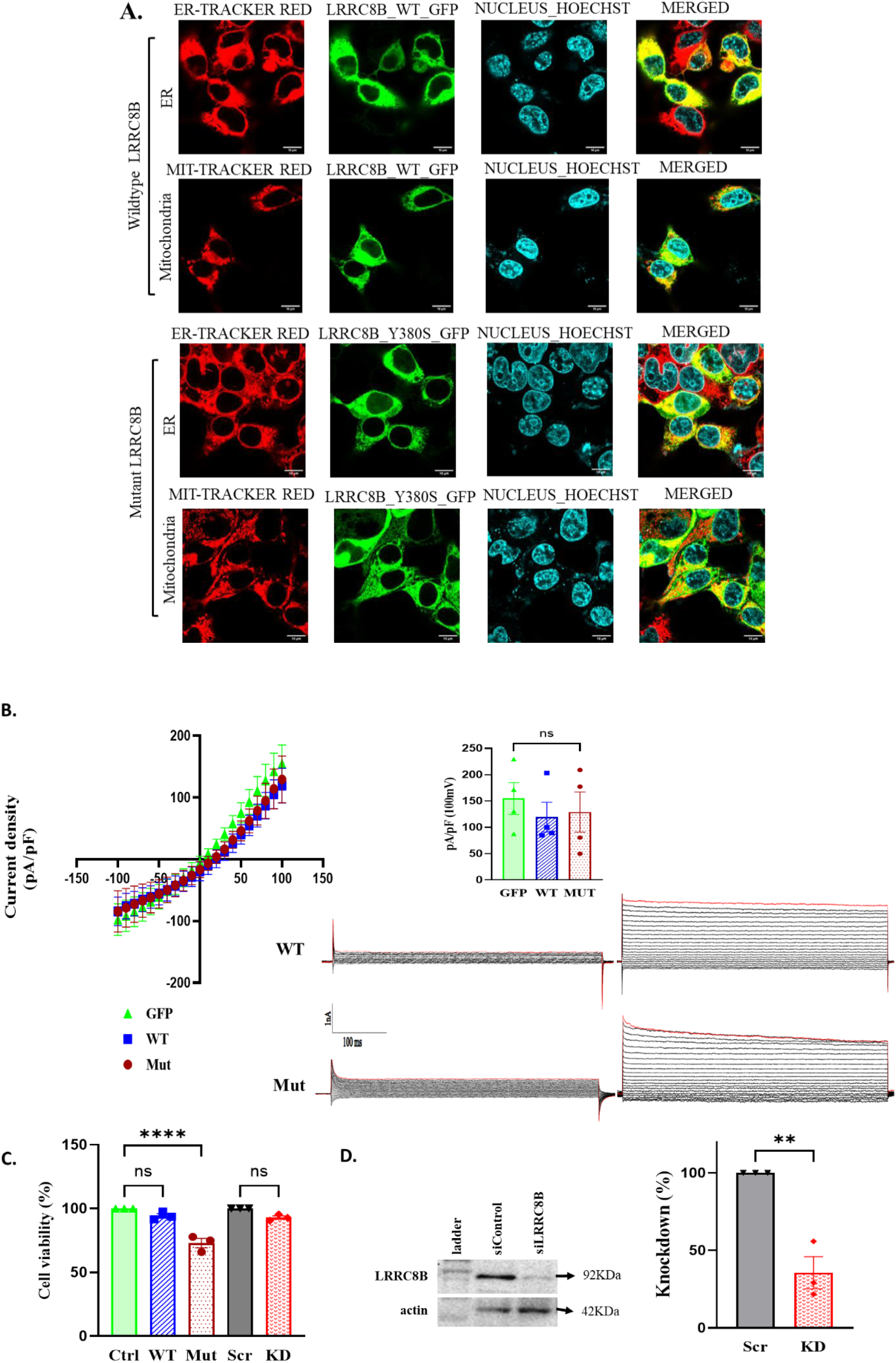
LRRC8B mutation does not alter its localization or VRAC activity but reduces cell viability. (A) Confocal images of HEK293T cells expressing GFP-LRRC8B (WT) or Y380S Mutant, co-stained with ER-Tracker, Mito-Tracker, and Hoechst. Merged images show ER and mitochondrial localization. Scale bar: 10 µm. (B) Whole-cell patch clamp analysis of current density–voltage relationships under isotonic and hypotonic conditions. No significant differences at +100 mV (mean ± SEM, N = 4; one-way ANOVA, ns). (C) Cell viability (MTT) in GFP(Ctrl), LRRC8B WT, LRRC8BY380S (Mut), siControl (Scr), and siLRRC8B (KD) cells (mean ± SEM, N = 3; one-way ANOVA, ****p < 0.0001). (D) LRRC8B knockdown validated by Western blot and densitometry normalized to actin (mean ± SEM, N = 3; unpaired t-test, **p < 0.01).

Since LRRC8B has not been shown to form functional VRAC, we next examined whether LRRC8B-Y380S affects endogenous VRAC activity. Overexpression of either wild-type or mutant LRRC8B had no effect on hypotonicity-induced VRAC currents in HEK293 cells (Fig. 1B).

To investigate the role of LRRC8B in cell survival, we silenced endogenous LRRC8B expression using siRNA. Western blot analysis confirmed effective knockdown, with a mean reduction of approximately 60% relative to scrambled control, quantified from three independent experiments (Fig. 1C and Fig. S4). MTT assays showed that overexpression of wild-type LRRC8B or siRNA-mediated knockdown did not affect cell viability. In contrast, expression of LRRC8B-Y380S significantly reduced cell viability in HEK293T cells, suggesting a deleterious effect that may contribute to disease manifestation (Fig. 1C).

### Mutant LRRC8B decreases the ER calcium leak and increases histamine-induced cytosolic Ca²⁺ rise

Having previously demonstrated that LRRC8B overexpression accelerates ER Ca²⁺ leak, we next examined whether the disease-associated mutation alters this function. ER Ca²⁺ levels were monitored using the genetically encoded indicator R-CEPIA1er following SERCA inhibition with 1 µM thapsigargin (TG) in nominally Ca²⁺-free extracellular solution. Cells overexpressing wild-type LRRC8B showed faster fluorescence decay than GFP controls, consistent with enhanced leak. In contrast, cells expressing the mutant LRRC8B displayed markedly slower decay, comparable to that of endogenous LRRC8B knockdown cells, despite the continued presence of endogenous protein (Fig. 2A). Leak kinetics were quantified as t_₁/₂_, the time to 50% reduction in normalized R-CEPIA1er fluorescence (Fig. 2B). Wild-type LRRC8B overexpression significantly shortened t_₁/₂_ relative to GFP controls, whereas endogenous LRRC8B knockdown prolonged it. Cells expressing mutant LRRC8B failed to reach 50% fluorescence decay within the 240 s recording window, achieving only ∼44% reduction, precluding reliable t_₁/₂_ estimation. Together, these data demonstrate that the LRRC8B mutation severely impairs ER Ca²⁺ leak.

**Figure 2.**
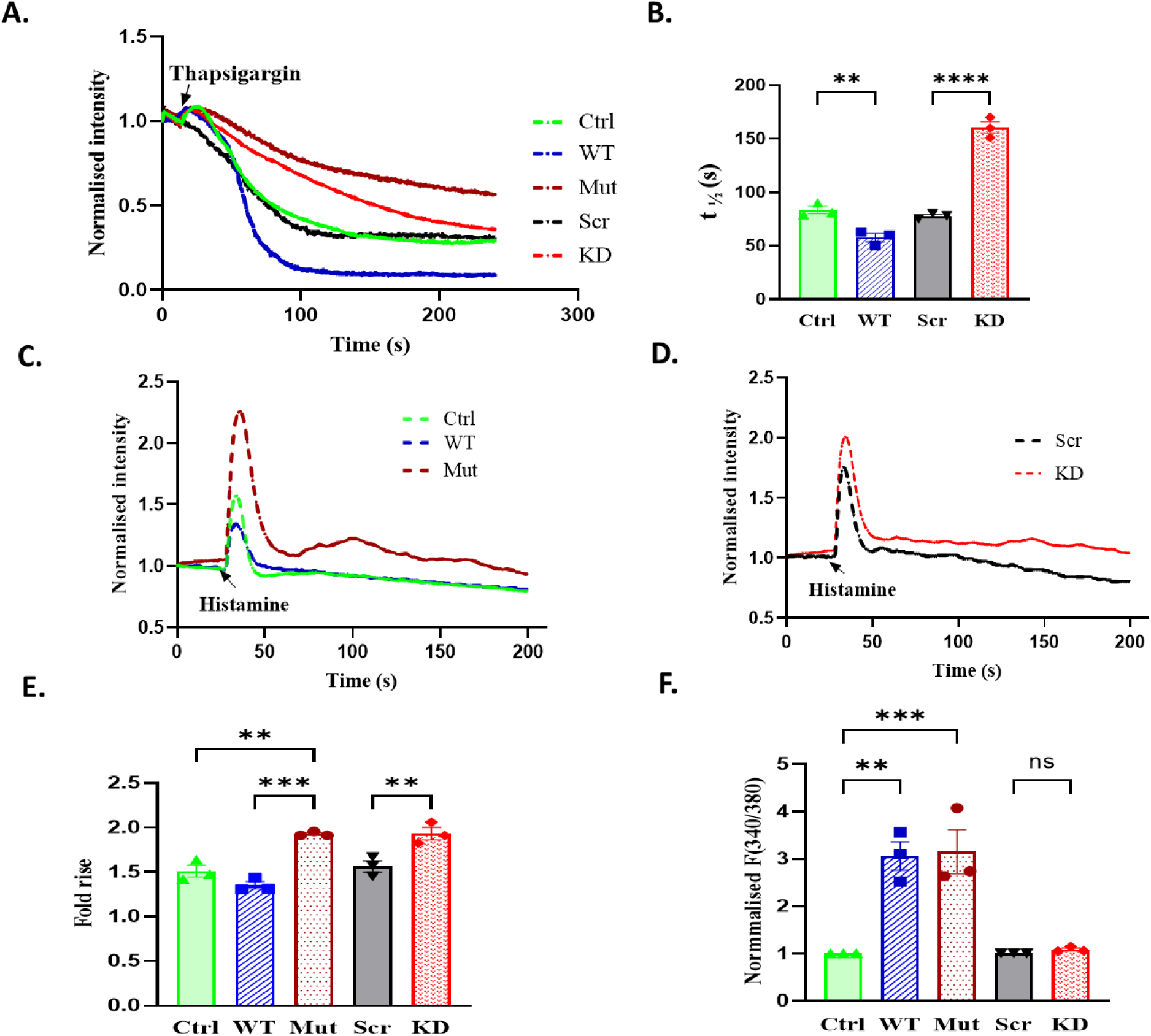
LRRC8B Mutant (Y380S) impairs ER Ca²⁺ leak, cytosolic Ca²⁺ dynamics, and steady-state cytosolic Ca²⁺ levels. (A) Representative traces of ER luminal Ca²⁺ decay after SERCA inhibition with 1 µM thapsigargin in nominally Ca²⁺-free solution, measured using R-CEPIA1er (normalized fluorescence). GFP(Ctrl) and scrambled siRNA (Scr) cells served as controls. (B) Quantification of t₁/₂ (time to 50% decay). LRRC8B Y380S (mut) cells did not reach t₁/₂ within 240 s (∼44% decay) and were excluded. Data are mean ± SEM (n = 15–30 cells, 3 independent experiments (N)); one-way ANOVA with Tukey’s test (**p < 0.01, ****p < 0.0001). (C–D) Representative CytoRCaMP traces after histamine stimulation in GFP (Ctrl), LRRC8B WT, LRRC8B Y380S (Mut) (C), and siControl (Scr) vs. siLRRC8B (KD) cells (D). Mutant cells show enhanced cytosolic Ca²⁺ peaks. (E) Peak cytosolic Ca²⁺ responses (fold change in CytoRCaMP). Mean ± SEM (n = 40–70 cells, N=4); one-way ANOVA with Tukey’s test (**p < 0.01, ***p < 0.001). (F) Basal cytosolic Ca²⁺ levels measured by fura-2 (340/380 ratio). Mean ± SEM (n = 30–80 cells, N=3); one-way ANOVA with Tukey’s test (**p < 0.01, ***p < 0.001).

To assess the functional consequences of impaired leak on ER Ca²⁺ handling, we measured the cytosolic Ca²⁺ rise ([Ca²⁺]_c_) evoked by histamine, which stimulates the Gq-coupled H1 receptor, triggering inositol 1,4,5-trisphosphate (IP₃) production and subsequent IP₃ receptor (IP₃R)-mediated ER Ca²⁺ release. Cytosolic Ca²⁺ was monitored using the genetically encoded indicator CytoRCaMP. Wild-type LRRC8B overexpression significantly attenuated the histamine-evoked [Ca²⁺]_c_ rise, consistent with reduced ER Ca²⁺ content resulting from enhanced constitutive leak (Fig. 2C). Conversely, endogenous LRRC8B knockdown augmented the [Ca²⁺]_c_ response (Fig. 2D). Strikingly, expression of mutant LRRC8B markedly enhanced IP₃R-mediated Ca²⁺ release and the subsequent [Ca²⁺]_c_ rise upon histamine stimulation, consistent with greater ER Ca²⁺ stores arising from impaired leak (Fig. 2E). Steady-state [Ca²⁺]_c_ levels were further assessed using the ratiometric dye Fura-2 AM. While LRRC8B knockdown had no measurable effect, overexpression of both wild-type and mutant LRRC8B significantly elevated steady-state [Ca²⁺]_c_ compared with controls (Fig. 2F).

### Effect of wild-type and mutant LRRC8B on mitochondrial Ca²⁺ uptake

Mitochondrial Ca²⁺ ([Ca²⁺]_m_) uptake was monitored using the mitochondria-targeted Ca²⁺ sensor R-CEPIA3mt. Histamine stimulation, which triggers IP₃-mediated ER Ca²⁺ release and subsequent mitochondrial Ca²⁺ uptake, revealed that overexpression of wild-type LRRC8B significantly enhanced the [Ca²⁺]_m_ rise, while knockdown of endogenous LRRC8B reduced it (Fig. 3A and 3B). Notably, expression of mutant LRRC8B markedly attenuated mitochondrial Ca²⁺ uptake compared to the respective control.

**Figure 3.**
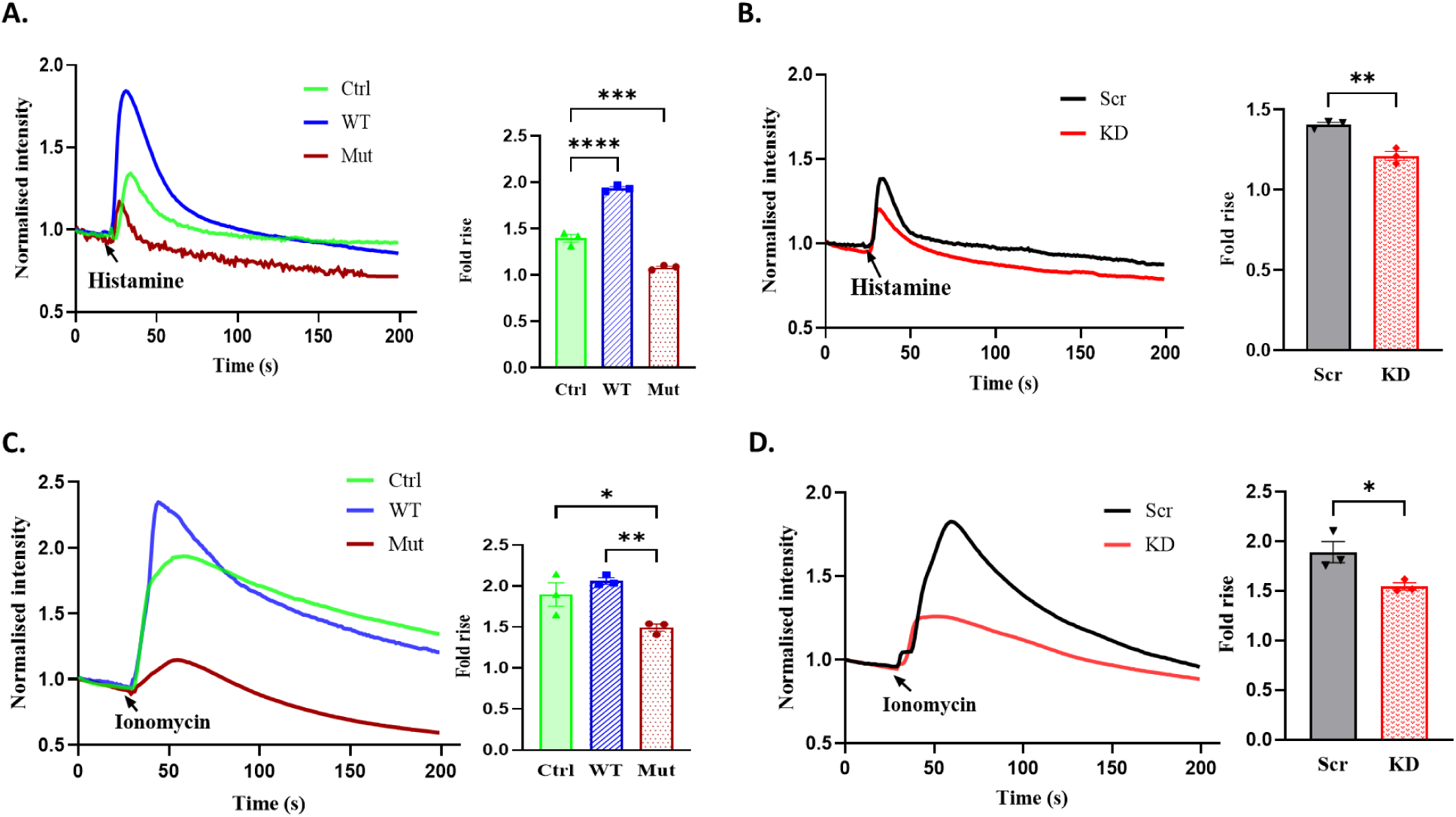
Effect of wild-type and mutant LRRC8B on mitochondrial Ca²⁺ uptake. (A) Representative traces of mitochondrial Ca²⁺ uptake (R-CEPIA3mt) following histamine stimulation in cells expressing GFP (Ctrl), LRRC8B WT, or LRRC8B Y380S (Mut), shown as normalized fluorescence over time. Bar graph depicts peak responses. Data are mean ± SEM (n = 12–30 cells per condition; N = 3). One-way ANOVA with Tukey’s test; **p < 0.01, ***p < 0.001. (B) Representative traces and quantification of mitochondrial Ca²⁺ uptake after histamine stimulation in cells treated with siControl (Scr) or siLRRC8B (KD). Data are mean ± SEM (n = 12–30 cells per condition; N = 3). Unpaired two-tailed Student’s t-test; **p < 0.01. (C) Representative traces and quantification of mitochondrial Ca²⁺ uptake following 5 µM ionomycin stimulation in cells expressing GFP (Ctrl), LRRC8B WT, or LRRC8B Y380S (Mut). Data are mean ± SEM (n = 30–50 cells per condition; N = 3). One-way ANOVA; *p < 0.05, **p < 0.01. (D) Representative traces and quantification of mitochondrial Ca²⁺ uptake after ionomycin stimulation in siControl (Scr) or siLRRC8B siRNA (KD)-treated cells. Data are mean ± SEM (n = 30–50 cells per condition; N = 3). Unpaired two-tailed Student’s t-test; **p < 0.01.

To determine whether this effect was specific to IP₃-mediated ER Ca²⁺ release, we elevated cytosolic [Ca²⁺]_c_ using a brief application of 5 µM ionomycin, a concentration that promotes Ca²⁺ entry across the plasma membrane (Yoshida et al., 2010). Ionomycin evoked a robust [Ca²⁺]_m_ rise; consistent with the histamine data, wild-type LRRC8B overexpression enhanced mitochondrial Ca²⁺ uptake, whereas both LRRC8B knockdown and mutant LRRC8B overexpression significantly reduced mitochondrial [Ca²⁺]_m_ rise compared to their respective controls (Fig. 3C and 3D). Together, these results demonstrate that LRRC8B positively regulates mitochondrial Ca²⁺ uptake, and that the disease-associated mutant acts in a dominant-negative manner, impairing mitochondrial Ca²⁺ handling even in the presence of endogenous LRRC8B.

### LRRC8B regulates mitochondrial membrane potential, ROS levels, and antioxidant gene expression

Given the role of LRRC8B in mitochondrial Ca²⁺ uptake, we examined its effect on mitochondrial membrane potential (ΔΨ_m_). Live-cell imaging with the potentiometric dye TMRE was used to assess ΔΨ_m_, with FCCP treatment serving as a control for complete depolarization (Fig. 4A–C). Application of FCCP results in a rapid drop in TMRE fluorescence, which is indicative of depolarization. The larger the fluorescence drop, the greater the ΔΨ_m_. Overexpression of LRRC8B-WT resulted in a significant increase in ΔΨ_m_ compared with vector controls, whereas LRRC8B knockdown had no measurable effect. In contrast, mutant LRRC8B overexpression led to a pronounced reduction in ΔΨ_m_, consistent with impaired mitochondrial health.

**Figure 4.**
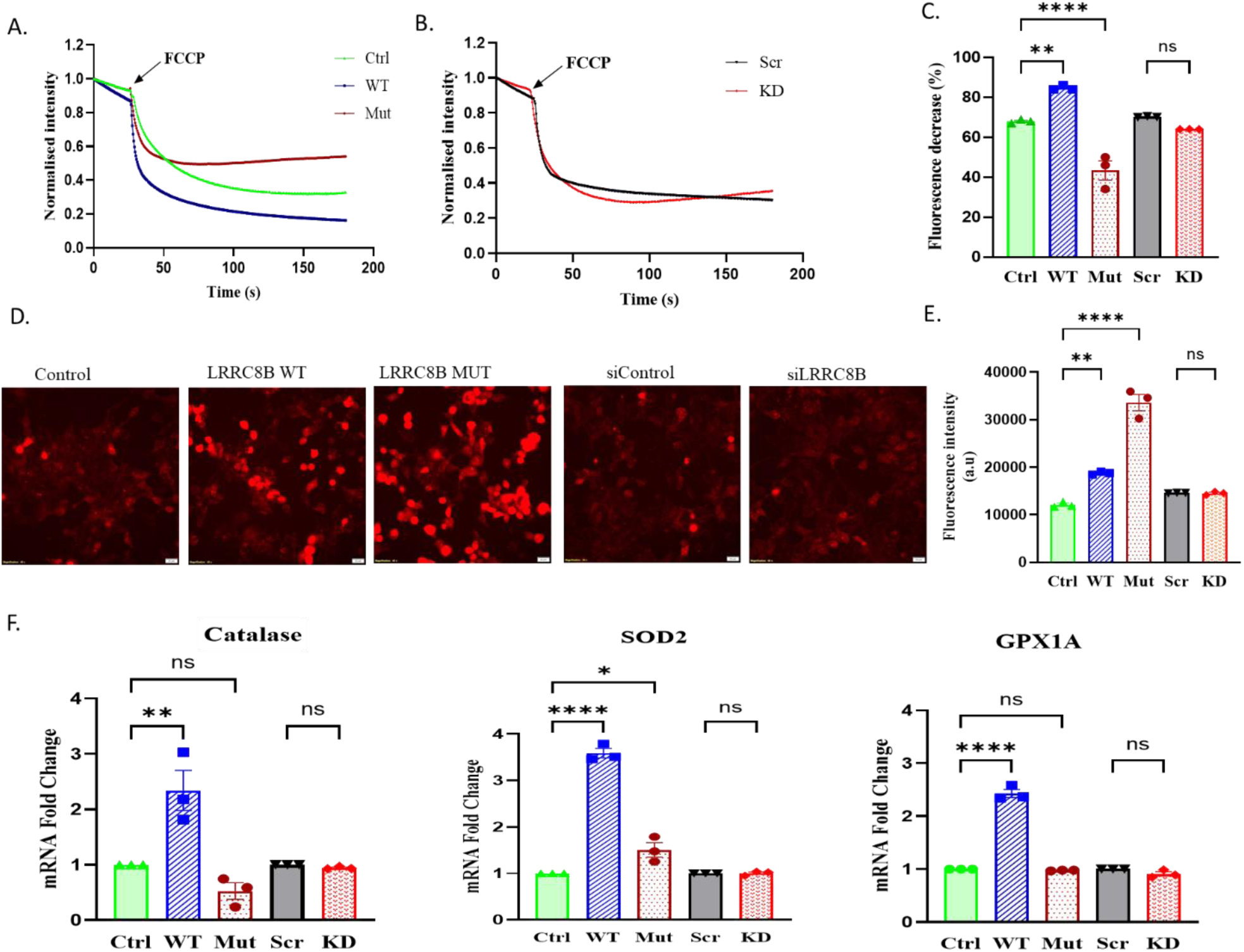
Effect of LRRC8B-WT and LRRC8B-Y380S on ΔΨ_m_, ROS, and antioxidant enzyme expression. (A–B) Representative TMRE traces showing mitochondrial membrane potential (ΔΨ_m_) after FCCP treatment in cells expressing (A) GFP (ctrl), LRRC8B wildtype (WT), or LRRC8B Y380S (Mut), and (B) siControl (Scr) or siLRRC8B (KD). (C) Percentage decrease in TMRE fluorescence following FCCP (mean ± SEM; n = 25–50 cells/condition, N = 3). (D) Representative MitoSOX images indicating mitochondrial superoxide levels in GFP (ctrl), LRRC8B WT, LRRC8B Y380S (Mut), siControl (Scr) and siLRRC8B (KD) cells. Scale bar, 20 μm. (E) Quantification of MitoSOX fluorescence (a.u.) (mean ± SEM; n = 30–100 cells/condition, N = 3). (F) RT–qPCR analysis of antioxidant genes (Catalase, SOD2, GPX1). LRRC8B WT significantly increased transcript levels compared to controls (mean ± SEM; N = 3). Statistics: One-way ANOVA with Tukey’s post hoc test; **p < 0.01, ****p < 0.0001; ns, not significant.

We then assessed mitochondrial oxidative stress by measuring superoxide levels using MitoSOX. Knockdown of LRRC8B did not alter mitochondrial superoxide levels, while both WT and mutant LRRC8B overexpression increased mitochondrial superoxide levels. Notably, mutant LRRC8B overexpression caused significantly higher superoxide accumulation than WT overexpression (Fig. 4D–E). To evaluate cellular antioxidant responses under these conditions, we measured transcript levels of superoxide dismutase 2 (SOD2), catalase, and glutathione peroxidase 1 (GPX1) using quantitative real-time PCR. In LRRC8B-WT–overexpressing cells, expression of all three antioxidant enzymes was upregulated, consistent with an adaptive response to elevated oxidative stress. In contrast, LRRC8B knockdown cells showed no change in antioxidant gene expression. Strikingly, mutant LRRC8B overexpression failed to induce catalase or GPX1 expression, despite elevated mitochondrial superoxide levels, suggesting an impaired antioxidant defense and increased susceptibility to oxidative stress (Fig. 4F).

### Effect of LRRC8B on mitochondrial metabolism

To assess the role of LRRC8B in mitochondrial metabolism, we measured oxygen consumption rate (OCR) using Seahorse analysis (Figure 5A-B). OCR was used as a readout of oxidative phosphorylation. Overexpression of LRRC8B-WT caused a small decrease in basal respiration and ATP-linked OCR. It also led to a slight increase in non-mitochondrial oxygen consumption. Maximal respiration was unchanged. LRRC8B knockdown did not affect basal respiration or ATP production. However, it increased non-mitochondrial oxygen consumption. It also reduced maximal respiratory capacity, indicating impaired reserve capacity. In contrast, cells expressing mutant LRRC8B showed a strong defect in mitochondrial function. Basal OCR, ATP production, and maximal respiration were all reduced. Non-mitochondrial oxygen consumption was markedly increased. These results suggest that LRRC8B-WT maintains normal mitochondrial function. Loss or mutation of LRRC8B impairs mitochondrial respiration. The mutant shows the strongest effect.

**Figure 5.**
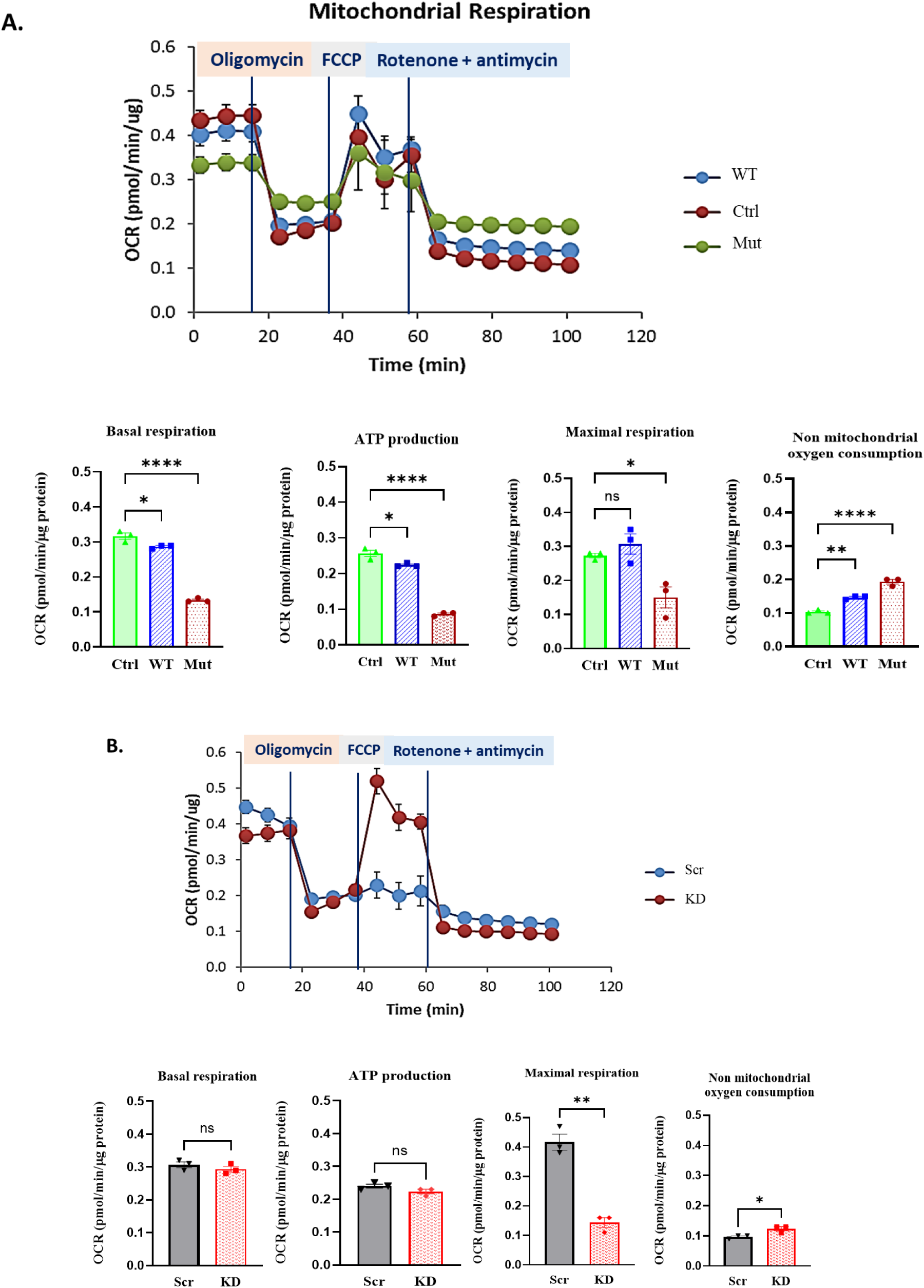
LRRC8B Regulates Mitochondrial Oxidative Phosphorylation. (A) OCR traces following sequential injections of oligomycin, FCCP, and rotenone/antimycin A in control(ctrl), LRRC8B wildtype (WT), and LRRC8B Y380S (Mut) expressing cells. Quantification shows that LRRC8B Mutant reduces basal respiration, ATP production, and maximal respiration, while increasing non-mitochondrial oxygen consumption. (B) OCR traces in siControl(scr) and siLRRC8B (KD) cells with corresponding quantification. Knockdown does not affect basal respiration or ATP production, but decreases maximal respiration and increases non-mitochondrial oxygen consumption. Data are mean ± SEM (N = 3). ns, not significant; *P < 0.05; **P < 0.01; ****P < 0.0001.

### LRRC8B mutation disrupts interaction with the mitochondrial VDAC

We investigated whether the Y380 mutation disrupts LRRC8B protein–protein interactions. Y380 is located in the C-terminus, proximal to the first leucine-rich repeat (LRR) domain (Fig. S1), and is conserved across all LRRC8 human paralogs and in different species (Fig. S2). Since the C-terminus of LRRC8 family members is cytoplasmic, it is accessible to cytosolic and organelle-associated proteins and mediates their interaction with LRRC8B. To identify interacting partners of LRRC8B, GFP-tagged wild-type and mutant LRRC8B were overexpressed in HEK293T cells, followed by GFP pull-down and LC–MS/MS analysis. Proteomic profiling revealed 12 proteins uniquely enriched in the WT interactome and 4 proteins selectively associated with the mutant (Fig. S3). Among the WT-specific interactors, the mitochondrial outer membrane protein voltage-dependent anion channel (VDAC) was identified, which was absent in the mutant interactome. VDAC is a key regulator of mitochondrial Ca²⁺ uptake, forming the primary conduit for Ca²⁺ entry across the outer mitochondrial membrane. It is enriched at mitochondria-associated membranes (MAMs), where it functionally couples ER-to-mitochondria Ca²⁺ transfer through interactions with IP₃R and the chaperone glucose-regulated protein 75 (GRP75), facilitating localized Ca²⁺ microdomain signaling. Given that VDAC is a direct mediator of mitochondrial Ca²⁺ entry, we validated this interaction by co-immunoprecipitation and immunoblotting.

Total VDAC protein levels were assessed in cells overexpressing LRRC8B-WT or LRRC8B-Y380S. Immunoblot analysis of total cell lysates and isolated mitochondrial fractions showed no significant difference in VDAC expression between conditions (Fig. 6A-C and Fig. S5). However, GFP pull-down followed by immunoblotting revealed a marked reduction in VDAC co-precipitation with LRRC8B-Y380S compared to WT (Fig. 6D-E and Fig. S6). Densitometric analysis confirmed a significant decrease in VDAC association with the mutant protein, despite comparable expression levels (Fig. 6F). Together, these findings indicate that the Y380S mutation selectively disrupts the LRRC8B–VDAC interaction without affecting VDAC abundance.

**Figure 6.**
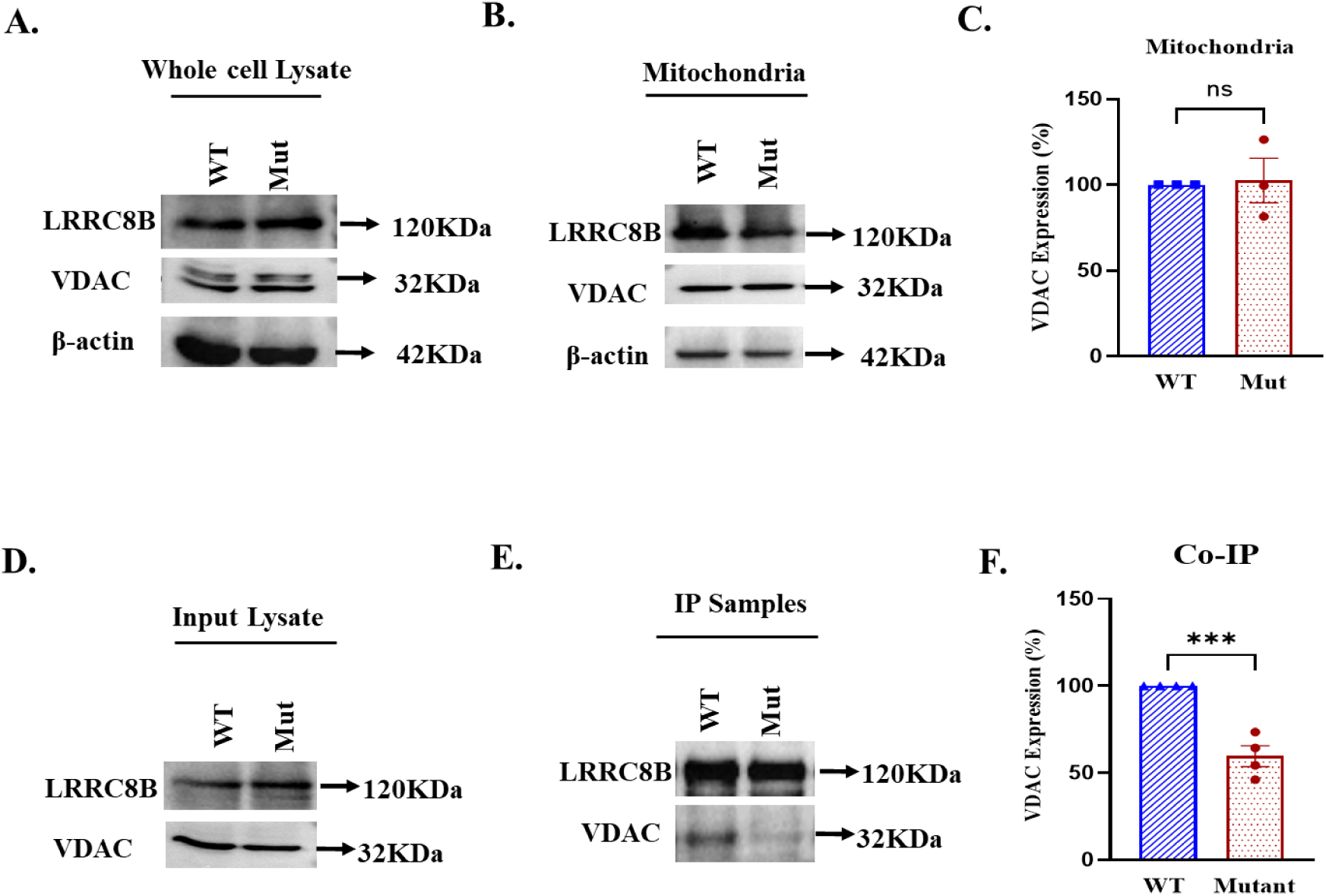
LRRC8B mutation selectively disrupts interaction with VDAC without altering its expression. (A–B) Immunoblot analysis of VDAC in whole-cell lysates (A) and mitochondrial fractions (B) from cells expressing LRRC8B wildtype (WT) or LRRC8B Y380S (Mut). (C) Densitometric quantification (normalized to actin) shows no difference in VDAC levels between conditions (mean ± SEM, N = 3; unpaired two-tailed t-test, ns). (D) Input blots confirm comparable expression of LRRC8B WT and LRRC8B Mutant. (E–F) Co-immunoprecipitation of LRRC8B-GFP (WT or mutant) followed by VDAC immunoblotting shows reduced VDAC association with LRRC8B Mutant. Quantification normalized to immunoprecipitated LRRC8B-GFP confirms this decrease (mean ± SEM, N = 3–4; unpaired two-tailed t-test, ***p < 0.001).

## Discussion

Disruption of intracellular Ca²⁺ homeostasis is a common pathogenic mechanism across neurological and psychiatric disorders. Pathogenic variants in voltage-gated Ca²⁺ channels (e.g., CACNA1C, CACNA1A), ligand-gated glutamate receptors, intracellular Ca²⁺ release channels (IP₃Rs, RyRs), and mitochondrial Ca²⁺ transport machinery impair Ca²⁺ flux, synaptic signaling, and neuronal bioenergetics (Filadi et al., 2017; Klocke et al., 2023; Ruiz et al., 2009; Szymanowicz et al., 2024; Zhou et al., 2022). Collectively, these findings establish dysregulated Ca²⁺ signaling as a unifying framework linking molecular defects to brain dysfunction. Within this framework, our findings position LRRC8B as a multifunctional Ca²⁺ regulatory protein operating at the intersection of ER Ca²⁺ homeostasis and mitochondrial Ca²⁺ uptake.

Previous work from our lab and others established LRRC8B as a regulator of ER Ca²⁺ leak (Ghosh et al., 2017) rather than a functional component of VRACs (Gaitán-Peñas et al., 2016). Consistent with this, LRRC8B suppression did not alter hypotonicity-induced VRAC currents, confirming its functionally distinct niche within the LRRC8 family. We investigated a disease-associated missense variant (Y380S) identified in patients with severe psychiatric disorders. Both wild-type and mutant LRRC8B localized to the ER and mitochondria, suggesting a role in ER-mitochondrial Ca²⁺ coupling. Wild-type LRRC8B overexpression enhanced ER Ca²⁺ leak, while knockdown reduced it. In contrast, Y380S overexpression markedly attenuated ER Ca²⁺ leak, phenocopying knockdown and resulting in increased ER Ca²⁺ content, evidenced by exaggerated histamine-evoked cytosolic Ca²⁺ transients. Paradoxically, mutant-expressing cells exhibited elevated steady-state cytosolic Ca²⁺ despite reduced ER leak, suggesting a possibility of impaired mitochondrial Ca²⁺ buffering capacity.

Ca²⁺ transfer from the ER to mitochondria occurs at mitochondria-associated ER membranes (MAMs), where the IP₃R-GRP75-VDAC complex generates high-amplitude Ca²⁺ microdomains sufficient to activate the low-affinity mitochondrial Ca²⁺ uniporter (MCU) (De Stefani et al., 2012; Rizzuto et al., 1993; Szabadkai et al., 2006). Given LRRC8B’s dual localization, we examined its role in mitochondrial Ca²⁺ uptake. Wild-type LRRC8B enhanced mitochondrial Ca²⁺ accumulation following IP₃-mediated ER Ca²⁺ release, while knockdown reduced uptake. Y380S severely impaired mitochondrial Ca²⁺ uptake regardless of endogenous LRRC8B levels, indicating a dominant-negative effect. We identified VDAC as a selective interactor of wild-type LRRC8B and showed that Y380S abolishes this interaction, providing a mechanistic basis for the observed phenotype. The Y380S substitution, located proximal to the first LRR domain, disrupts the LRRC8B-VDAC interface without affecting VDAC expression or mitochondrial abundance. Consequently, loss of this coupling reduces the efficiency of ER-derived Ca²⁺ microdomains reaching the outer mitochondrial membrane (OMM), attenuating MCU activation.

Beyond ER-mitochondria coupling, ionomycin experiments reveal that wild-type LRRC8B also enhances mitochondrial Ca²⁺ uptake independently of IP₃R-mediated ER release. At 5 µM ionomycin, that makes plasma membrane permeable to Ca²⁺, wild-type LRRC8B overexpression still enhanced mitochondrial Ca²⁺ uptake, whereas knockdown and Y380S attenuated it. This suggests LRRC8B may facilitate mitochondrial Ca²⁺ uptake through mechanisms that include modulation of OMM Ca²⁺ permeability via VDAC interaction or maintenance of mitochondrial membrane potential (ΔΨ_m_). Our TMRE data confirm that wild-type LRRC8B overexpression significantly enhances ΔΨ_m_, the primary electrochemical driving force for MCU-mediated Ca²⁺ import, thereby amplifying mitochondrial Ca²⁺ uptake capacity across diverse Ca²⁺ delivery routes. An intriguing, though speculative, possibility is that LRRC8B may contribute to a Ca²⁺-permeable pathway on the mitochondrial membrane, given its four transmembrane domains shared with pore-forming LRRC8 family members and the emerging recognition that LRRC8 proteins can assemble into functional channels beyond the plasma membrane, including in intracellular organelles such as lysosomes (Li et al., 2020).

The functional consequences of impaired LRRC8B-VDAC interaction are manifested across multiple indices of mitochondrial health. Mitochondrial Ca²⁺ is an obligate activator of key TCA cycle dehydrogenases driving NADH production and oxidative phosphorylation (OXPHOS) (Denton, 2009; Denton & McCormack, 1986). Consistent with this, Y380S expression caused significant reductions in basal OCR, ATP-linked respiration, and maximal respiratory capacity, accompanied by ΔΨ_m_ collapse, confirming severe OXPHOS failure. By contrast, wild-type LRRC8B overexpression preserved maximal respiratory capacity and modestly elevated ΔΨ_m_. Knockdown cells maintained basal bioenergetics while showing diminished maximal respiratory capacity, suggesting residual mitochondrial Ca²⁺ uptake is sufficient for baseline OXPHOS under non-stressed conditions.

The marked elevation of mitochondrial superoxide in Y380S-expressing cells is concordant with severe OXPHOS dysfunction, as electron leak from a compromised respiratory chain generates superoxide as a major by-product. Notably, wild-type LRRC8B overexpression, despite modestly increasing mitochondrial superoxide, induced coordinated transcriptional upregulation of antioxidant enzymes SOD2, catalase, and GPX1, reflecting a physiologically adaptive mitohormesis response (Ristow & Schmeisser, 2014). Y380S-expressing cells failed to mount this adaptive response despite markedly higher superoxide levels, indicating a breakdown in mitochondrial retrograde signaling, the pathway by which mitochondrial status is communicated to the nucleus to orchestrate adaptive gene expression (Butow & Avadhani, 2004).

The reduced cell viability seen with Y380S overexpression, but not with knockdown or wild-type overexpression, suggests that the mutant actively disrupts cell survival rather than simply causing a loss of LRRC8B function. This likely reflects a combination of dominant-negative impairment of ER Ca²⁺ leak, disrupted LRRC8B-VDAC interaction, collapse of mitochondrial bioenergetics, and failure of antioxidant defenses, a pleiotropic cellular catastrophe that compensatory mechanisms cannot fully offset. The postmitotic nature of neurons, their exceptionally high energy demands, and critical dependence on OXPHOS (Harris et al., 2012; Rumpf et al., 2023) render them particularly vulnerable to such mitochondrial dysfunction. Given the high expression of LRRC8B in the brain (Pervaiz et al., 2019), any mutation-driven impairment of its function could affect neuronal survival and activity. Together, these findings provide a mechanistic basis for how the Y380S variant, identified in patients with severe mental illness (Ganesh et al., 2019), could impair neuronal Ca²⁺ homeostasis, bioenergetics, and survival.

## Limitations

We acknowledge several limitations of the present study and are actively working to address them. First, all experiments were performed in HEK293T cells. While this system is widely used for Ca²⁺ signaling and mitochondrial studies, it does not fully capture the physiological context of neurons or other brain cell types in which the Y380S variant is expected to exert its pathogenic effects. To address this, we are currently evaluating whether the observed phenotypes are conserved in iPSC-derived neurons, primary neuronal cultures, and relevant brain organoid models.

Second, although LRRC8B was found to localize to both the ER and mitochondria, we did not directly assess its enrichment at MAMs or its role in MAM formation and integrity. These aspects are under active investigation.

Third, we identified VDAC as a selective interactor of wild-type LRRC8B. However, the specific isoform(s) involved, VDAC1, VDAC2, or VDAC3, remain to be defined. In addition, whether this interaction is direct or mediated by intermediary proteins such as GRP75 or IP₃R has not yet been resolved. Ongoing experiments are aimed at clarifying the molecular nature of this interaction.

Fourth, the electrophysiological basis of LRRC8B-mediated ER Ca²⁺ leak was not directly examined. The mechanism by which LRRC8B facilitates Ca²⁺ efflux, whether as a pore-forming component, a channel modulator, or an accessory regulator of established leak pathways, such as TRIC channels, pannexin-1, or the Sec61 translocon, will require direct electrophysiological measurements and reconstitution approaches. These studies are currently in progress.

Fifth, the possibility that LRRC8B forms a Ca²⁺-permeable channel in the mitochondrial membrane remains a working hypothesis that has not yet been tested experimentally. We are pursuing mitoplast patch-clamp recordings, Ca²⁺ flux assays in isolated mitochondria, and proteoliposome reconstitution with purified LRRC8B to rigorously evaluate this model. Finally, this study relied on acute overexpression and siRNA-mediated knockdown approaches, which may not fully recapitulate the chronic heterozygous expression of the Y380S variant observed in patients. To overcome this limitation, we are developing knock-in models that express the variant at endogenous levels, enabling more physiologically relevant validation.

## Acknowledgement

We acknowledge Rajamathanki and Dr. Gomathy for thier critical comments on the manuscript.

## Author contribution statement

Conceptualization: AA and AKB. Methodology, investigation, Formal analysis: AA, DS, RS, AG. Writing (original draft, review & editing): AA, DS, and AKB. Supervision; Project, Funding acquisition: AKB.

## Declaration of generative AI and AI-assisted technologies in the writing process

During the preparation of this work, the authors used ChatGPT/Claude in order to improve readability and language. After using this tool, the authors reviewed and edited the content as needed and take full responsibility for the content of the publication. Graphical Abstract and schematic diagrams are created using Biorender software.

## Data availability statement

The data used to support the findings of this study are available from the corresponding author upon request.

## Supplementary Data

**Figure S1.**
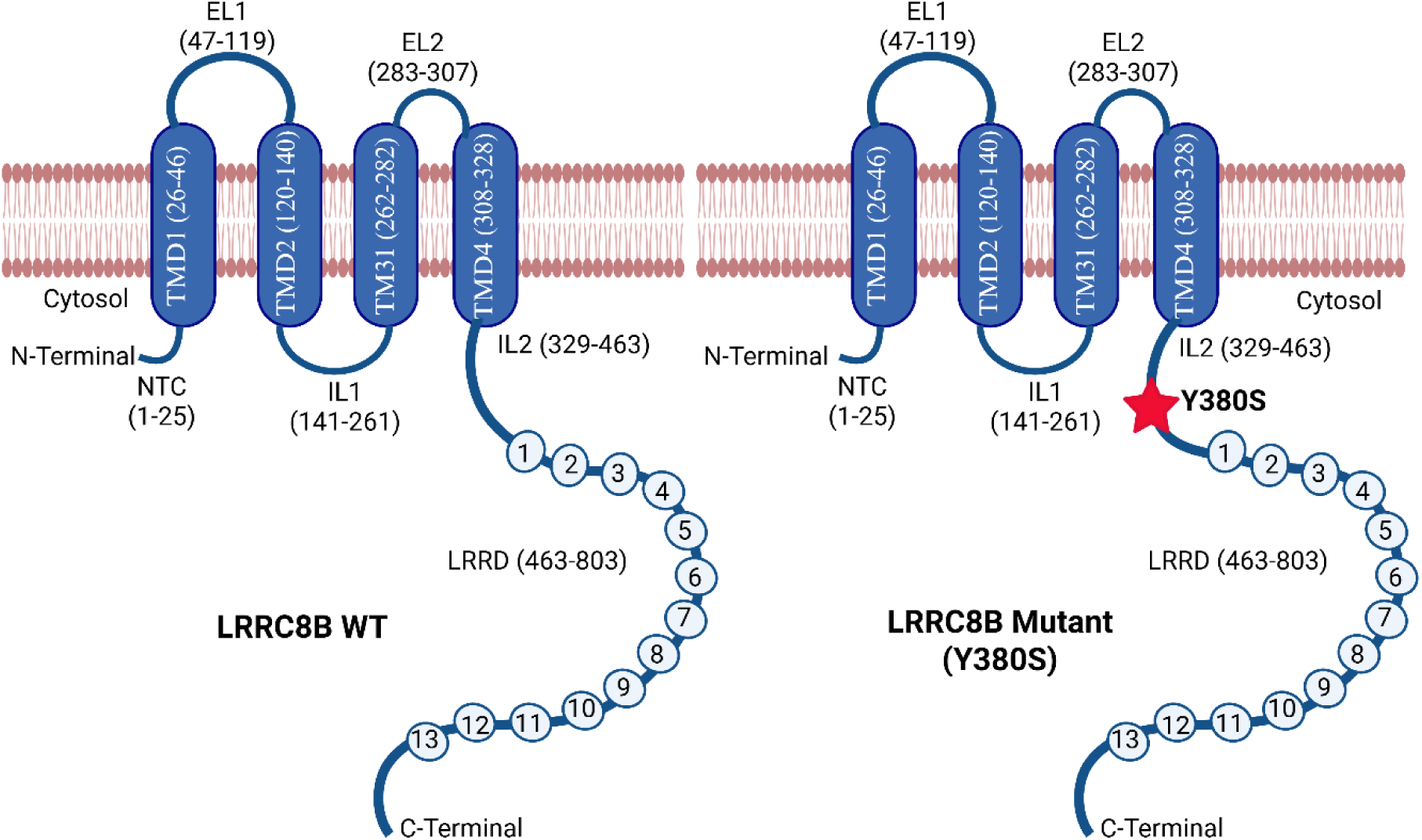
Schematic representation of wild-type and mutant LRRC8B topology. Diagram illustrating the membrane topology of LRRC8B, comprising 803 amino acids and four transmembrane domains (TM1: residues 26–46; TM2: 120–140; TM3: 262–282; TM4: 308–328). Both N- and C-termini are oriented toward the cytosol. The Y380S mutation (indicated by a red star) is located in the cytosolic region, proximal to the leucine-rich repeat domain (LRRD; residues 463–803), which contains 13 leucine-rich repeats. (Created in https://BioRender.com)

**Figure S2.**
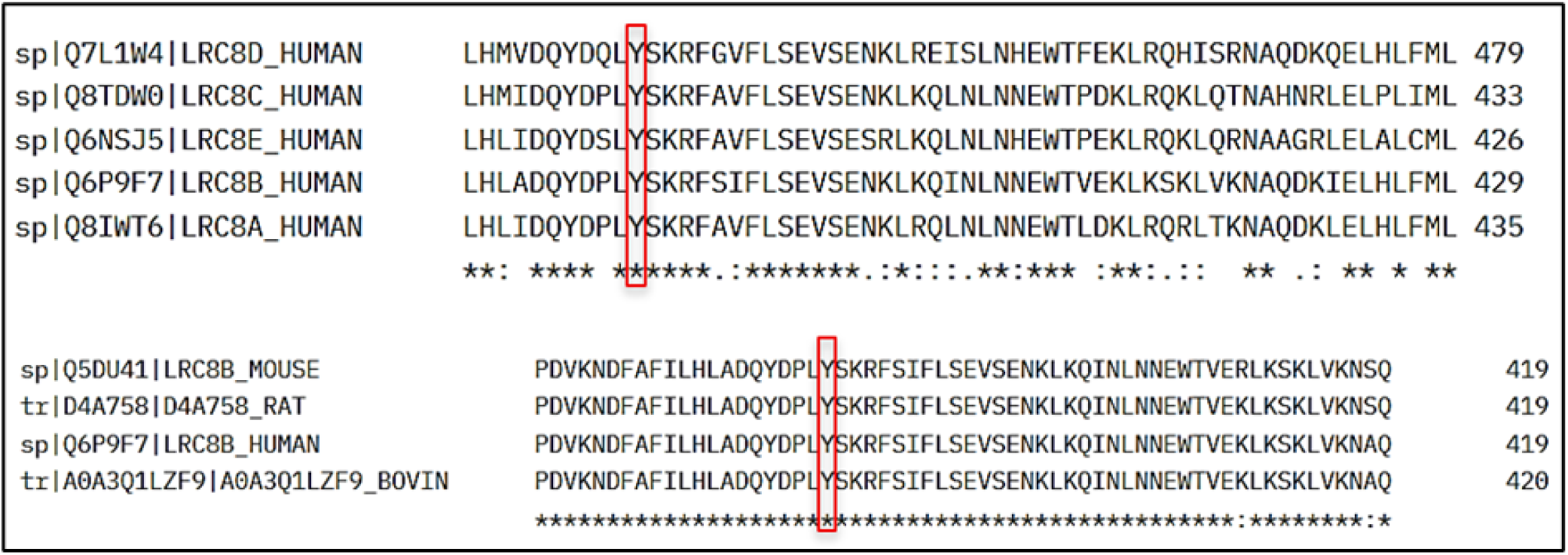
Conservation of tyrosine (Y) at position 380 in LRRC8B. Multiple sequence alignment showing conservation of the tyrosine residue (Y380) in LRRC8B across human LRRC8 paralogs (LRRC8A, LRRC8C, LRRC8D, and LRRC8E) and among different species, including human, mouse, rat, and bovine. The conserved tyrosine residue is highlighted in red. Asterisks indicate fully conserved residues across sequences.

**Figure S3.**
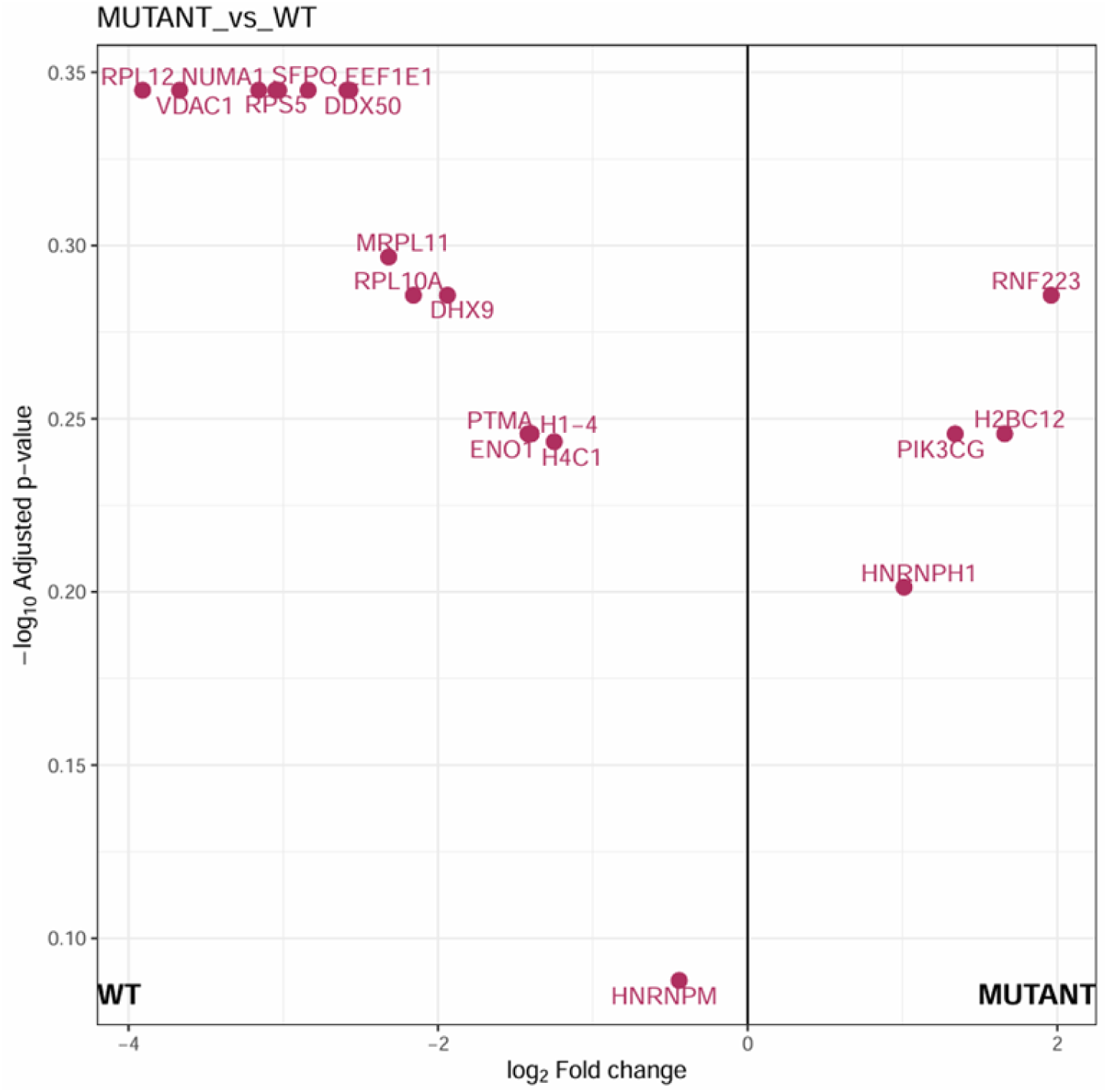
Differential interactome analysis of LRRC8B WT and mutant. Volcano plot showing LC–MS/MS–based proteomic analysis of GFP pull-down samples from HEK293T cells expressing GFP-tagged wild-type (WT) or mutant LRRC8B. The x-axis represents log₂ fold change (mutant vs WT), and the y-axis shows −log₁₀ adjusted p-value. Proteins enriched in WT samples are shown on the left, while those enriched in mutant samples are shown on the right. Selected significantly enriched proteins are labeled. VDAC1 is identified as a WT-enriched interactor.

**Figure S4.**
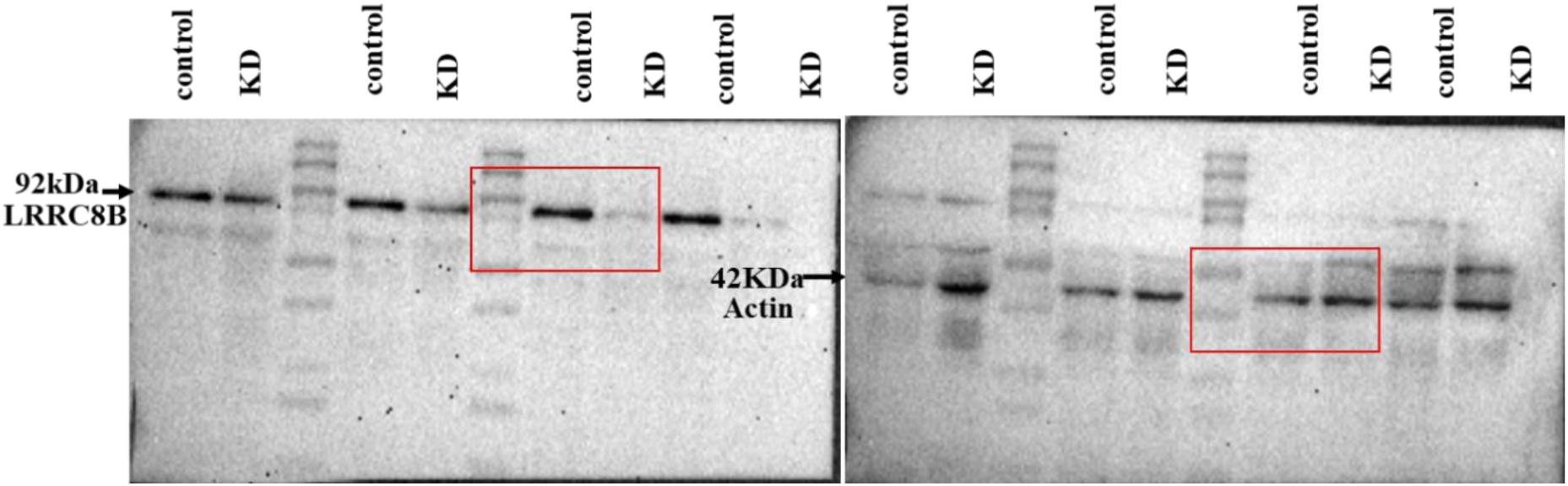
Full-length immunoblot images for siRNA validation. Uncropped western blot images corresponding to the siRNA-mediated knockdown experiments shown in the main figures. Cells were transfected with control siRNA or siLRRC8B (KD), and protein lysates were analyzed by immunoblotting. The red boxes indicate the regions that were cropped and presented in the main text figures. Actin (∼42 kDa) is shown as a loading control. Molecular weight markers are indicated where applicable.

**Figure S5.**
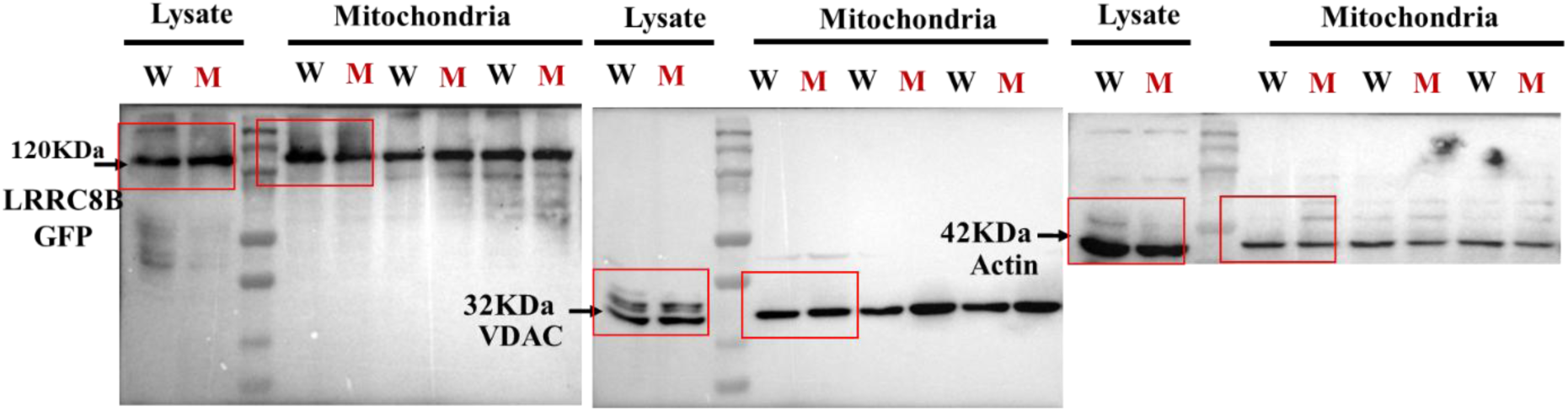
Quantification of VDAC protein expression levels. Representative immunoblot images showing VDAC protein levels in total cell lysates and isolated mitochondrial fractions from cells transfected with GFP-tagged LRRC8B WT (W) or mutant Y380S (M) constructs. Actin (∼42 kDa) is included as a loading control for total lysates. Red boxes indicate the regions that were cropped and presented in the main figures. Molecular weight markers are shown where applicable.

**Figure S6.**
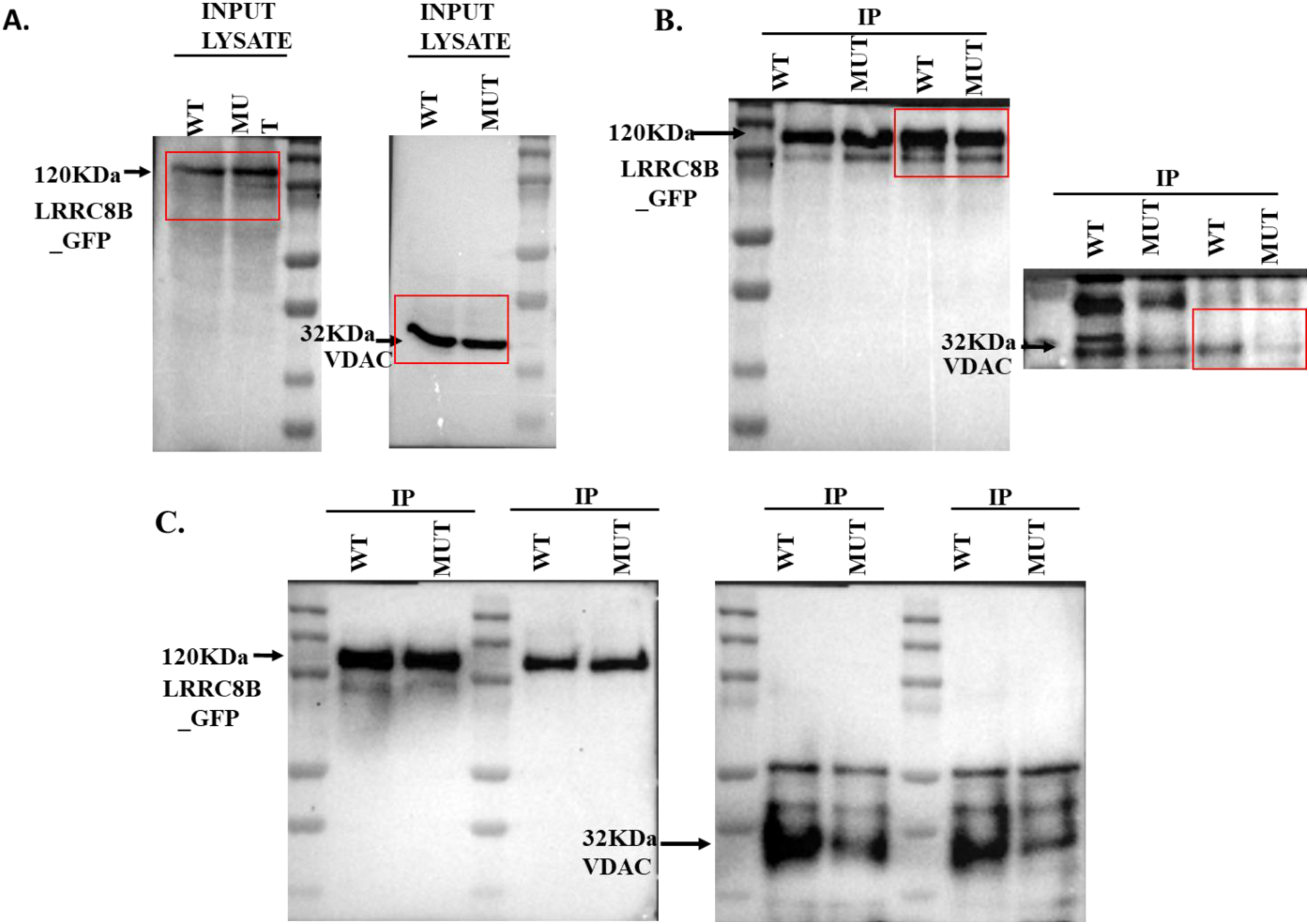
VDAC protein levels and co-immunoprecipitation with LRRC8B. **(A)** Representative immunoblot images showing protein levels of GFP-tagged LRRC8B (WT and mutant) and VDAC in input lysates used for pull-down assays. **(B–C)** Immunoblot analysis of four independent immunoprecipitation (IP) experiments. Bands at ∼120 kDa confirm successful pull-down of GFP-tagged LRRC8B (WT and mutant) using an anti-GFP antibody. VDAC (∼32 kDa) bands indicate co-precipitation of VDAC with LRRC8B WT and mutant proteins. VDAC levels were normalized to the corresponding LRRC8B (WT or mutant) levels for quantification. Red boxes highlight the regions that were cropped and presented in the main figures. Molecular weight markers are shown where applicable.

**Table S1.**
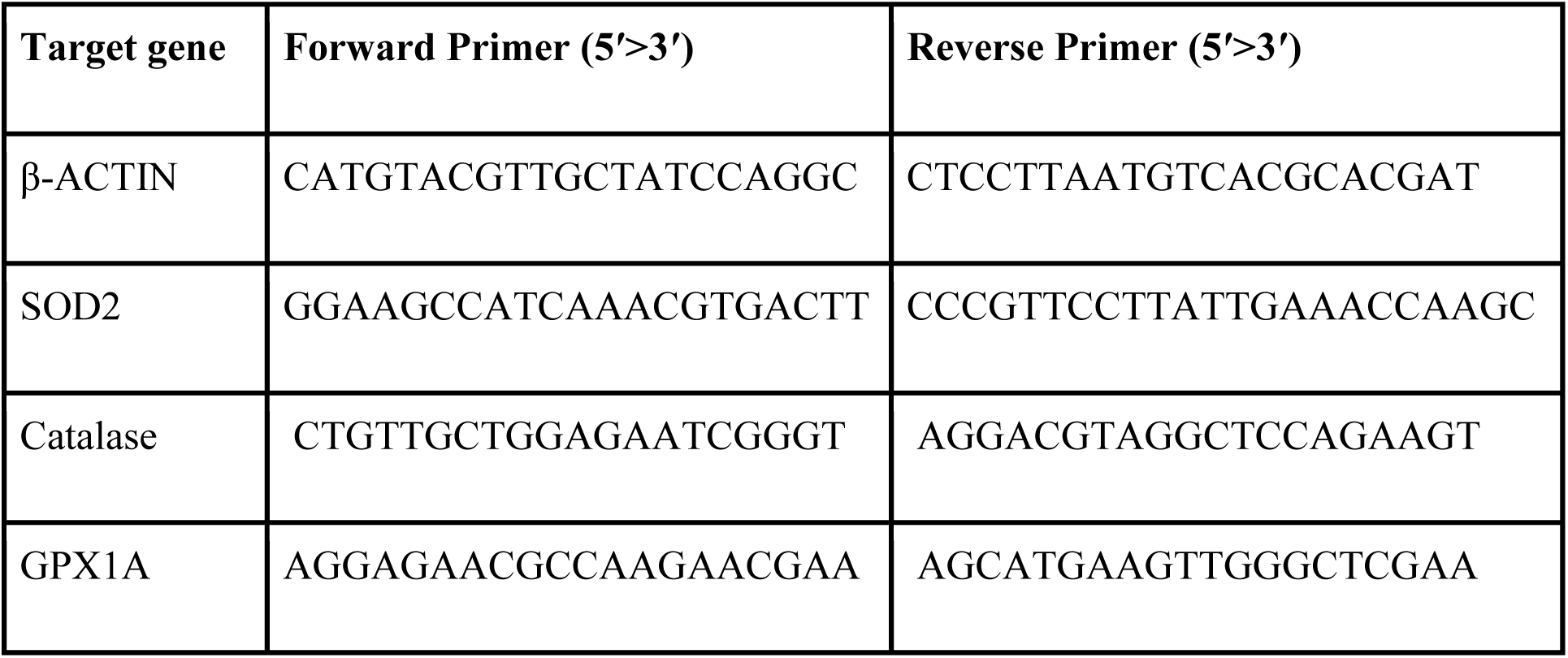
List of RT-PCR primers used in this study.

## References

Abascal, F., & Zardoya, R. (2012). LRRC8 proteins share a common ancestor with pannexins, and may form hexameric channels involved in cell-cell communication. *BioEssays: News and Reviews in Molecular*, Cellular and Developmental Biology, 34(7), 551–560. 10.1002/bies.201100173

Butow, R. A., & Avadhani, N. G. (2004). Mitochondrial Signaling: The Retrograde Response. Molecular Cell, 14(1), 1–15. 10.1016/S1097-2765(04)00179-0

De Stefani, D., Bononi, A., Romagnoli, A., Messina, A., De Pinto, V., Pinton, P., & Rizzuto, R. (2012). VDAC1 selectively transfers apoptotic Ca2+ signals to mitochondria. Cell Death & Differentiation, 19(2), 267–273. 10.1038/cdd.2011.92

Denton, R. M. (2009). Regulation of mitochondrial dehydrogenases by calcium ions. *Biochimica et Biophysica Acta (BBA) - Bioenergetics*, Mitochondrial Calcium in Health and Disease, 1787(11), 1309–1316. 10.1016/j.bbabio.2009.01.005

Denton, R. M., & McCormack, J. G. (1986). The calcium sensitive dehydrogenases of vertebrate mitochondria. Cell Calcium, 7(5), 377–386. 10.1016/0143-4160(86)90040-0

Filadi, R., Theurey, P., & Pizzo, P. (2017). The endoplasmic reticulum-mitochondria coupling in health and disease: Molecules, functions and significance. Cell Calcium, 62, 1–15. 10.1016/j.ceca.2017.01.003

Gaitán-Peñas, H., Gradogna, A., Laparra-Cuervo, L., Solsona, C., Fernández-Dueñas, V., Barrallo-Gimeno, A., Ciruela, F., Lakadamyali, M., Pusch, M., & Estévez, R. (2016). Investigation of LRRC8-Mediated Volume-Regulated Anion Currents in Xenopus Oocytes. Biophysical Journal, 111(7), 1429–1443. 10.1016/j.bpj.2016.08.030

Ganesh, S., Ahmed P., H., Nadella, R. K., More, R. P., Seshadri, M., Viswanath, B., Rao, M., Jain, S., Consortium, T. A., & Mukherjee, O. (2019). Exome sequencing in families with severe mental illness identifies novel and rare variants in genes implicated in Mendelian neuropsychiatric syndromes. Psychiatry and Clinical Neurosciences, 73(1), 11–19. 10.1111/pcn.12788

Ghosh, A., Khandelwal, N., Kumar, A., & Bera, A. K. (2017). Leucine-rich repeat-containing 8B protein is associated with the endoplasmic reticulum Ca2+ leak in HEK293 cells. Journal of Cell Science, 130(22), 3818–3828. 10.1242/jcs.203646

Harris, J. J., Jolivet, R., & Attwell, D. (2012). Synaptic Energy Use and Supply. Neuron, 75(5), 762–777. 10.1016/j.neuron.2012.08.019

Hirabayashi, Y., Kwon, S.-K., Paek, H., Pernice, W. M., Paul, M. A., Lee, J., Erfani, P., Raczkowski, A., Petrey, D. S., Pon, L. A., & Polleux, F. (2017). ER-mitochondria tethering by PDZD8 regulates Ca2+ dynamics in mammalian neurons. Science, 358(6363), 623–630. 10.1126/science.aan6009

Huang, C.-Y., Chiang, S.-F., Lin, T.-Y., Chiou, S.-H., & Chow, K.-C. (2012). HIV-1 Vpr Triggers Mitochondrial Destruction by Impairing Mfn2-Mediated ER-Mitochondria Interaction. PLOS ONE, 7(3), e33657. 10.1371/journal.pone.0033657

Kanemaru, K., Suzuki, J., Taiko, I., & Iino, M. (2020). Red fluorescent CEPIA indicators for visualization of Ca2+ dynamics in mitochondria. Scientific Reports, 10(1), 2835. 10.1038/s41598-020-59707-8

Karakas, E., Strange, K., & Denton, J. S. (2025). Recent advances in structural characterization of volume-regulated anion channels (VRACs). The Journal of Physiology, 603(15), 4201–4211. 10.1113/JP286189

Klocke, B., Krone, K., Tornes, J., Moore, C., Ott, H., & Pitychoutis, P. M. (2023). Insights into the role of intracellular calcium signaling in the neurobiology of neurodevelopmental disorders. Frontiers in Neuroscience, 17. 10.3389/fnins.2023.1093099

Kubota, K., Kim, J. Y., Sawada, A., Tokimasa, S., Fujisaki, H., Matsuda-Hashii, Y., Ozono, K., & Hara, J. (2004). LRRC8 involved in B cell development belongs to a novel family of leucine-rich repeat proteins. FEBS Letters, 564(1–2), 147–152. 10.1016/S0014-5793(04)00332-1

Li, P., Hu, M., Wang, C., Feng, X., Zhao, Z., Yang, Y., Sahoo, N., Gu, M., Yang, Y., Xiao, S., Sah, R., Cover, T. L., Chou, J., Geha, R., Benavides, F., Hume, R. I., & Xu, H. (2020). LRRC8 family proteins within lysosomes regulate cellular osmoregulation and enhance cell survival to multiple physiological stresses. Proceedings of the National Academy of Sciences, 117(46), 29155–29165. 10.1073/pnas.2016539117

Lutter, D., Ullrich, F., Lueck, J. C., Kempa, S., & Jentsch, T. J. (2017). Selective transport of neurotransmitters and modulators by distinct volume-regulated LRRC8 anion channels. Journal of Cell Science, 130(6), 1122–1133. 10.1242/jcs.196253

Marchi, S., Patergnani, S., & Pinton, P. (2014). The endoplasmic reticulum-mitochondria connection: One touch, multiple functions. Biochimica Et Biophysica Acta, 1837(4), 461–469. 10.1016/j.bbabio.2013.10.015

Miyake, K., Yamashita, Y., Ogata, M., Sudo, T., & Kimoto, M. (1995). RP105, a novel B cell surface molecule implicated in B cell activation, is a member of the leucine-rich repeat protein family. Journal of Immunology (Baltimore, Md.: 1950), 154(7), 3333–3340.

Mustaly-Kalimi, S., Gallegos, W., Steinbrenner, D., Gupta, S., Houcek, A. J., Bennett, D. A., Marr, R. A., Peterson, D. A., Sekler, I., & Stutzmann, G. E. (2025). Mitochondrial dysfunction mediated by ER-calcium dysregulation in neurons derived from Alzheimer’s disease patients. Acta Neuropathologica Communications, 13, 165. 10.1186/s40478-025-02023-x

Pervaiz, S., Kopp, A., von Kleist, L., & Stauber, T. (2019). Absolute Protein Amounts and Relative Abundance of Volume-regulated Anion Channel (VRAC) LRRC8 Subunits in Cells and Tissues Revealed by Quantitative Immunoblotting. International Journal of Molecular Sciences, 20(23), 5879. 10.3390/ijms20235879

Qiu, Z., Dubin, A. E., Mathur, J., Tu, B., Reddy, K., Miraglia, L. J., Reinhardt, J., Orth, A. P., & Patapoutian, A. (2014). SWELL1, a Plasma Membrane Protein, Is an Essential Component of Volume-Regulated Anion Channel. Cell, 157(2), 447–458. 10.1016/j.cell.2014.03.024

Ristow, M., & Schmeisser, K. (2014). Mitohormesis: Promoting Health and Lifespan by Increased Levels of Reactive Oxygen Species (ROS). Dose-Response, 12(2), dose-response.13-035.Ristow. 10.2203/dose-response.13-035.Ristow

Rizzuto, R., Brini, M., Murgia, M., & Pozzan, T. (1993). Microdomains with high Ca2+ close to IP3-sensitive channels that are sensed by neighboring mitochondria. Science, 262(5134), 744–747. 10.1126/science.8235595

Rizzuto, R., De Stefani, D., Raffaello, A., & Mammucari, C. (2012). Mitochondria as sensors and regulators of calcium signalling. Nature Reviews Molecular Cell Biology, 13(9), 566–578. 10.1038/nrm3412

Ruiz, A., Matute, C., & Alberdi, E. (2009). Endoplasmic reticulum Ca(2+) release through ryanodine and IP(3) receptors contributes to neuronal excitotoxicity. Cell Calcium, 46(4), 273–281. 10.1016/j.ceca.2009.08.005

Rumpf, S., Sanal, N., & Marzano, M. (2023). Energy metabolic pathways in neuronal development and function. Oxford Open Neuroscience, 2, kvad004. 10.1093/oons/kvad004

Sawada, A., Takihara, Y., Kim, J. Y., Matsuda-Hashii, Y., Tokimasa, S., Fujisaki, H., Kubota, K., Endo, H., Onodera, T., Ohta, H., Ozono, K., & Hara, J. (2003). A congenital mutation of the novel gene LRRC8 causes agammaglobulinemia in humans. Journal of Clinical Investigation, 112(11), 1707–1713. 10.1172/JCI18937

Schober, A. L., Wilson, C. S., & Mongin, A. A. (2017). Molecular composition and heterogeneity of the LRRC8-containing swelling-activated osmolyte channels in primary rat astrocytes. The Journal of Physiology, 595(22), 6939. 10.1113/JP275053

Suzuki, J., Kanemaru, K., Ishii, K., Ohkura, M., Okubo, Y., & Iino, M. (2014). Imaging intraorganellar Ca2+ at subcellular resolution using CEPIA. Nature Communications, 5(1), 4153. 10.1038/ncomms5153

Syeda, R., Qiu, Z., Dubin, A. E., Murthy, S. E., Florendo, M. N., Mason, D. E., Mathur, J., Cahalan, S. M., Peters, E. C., Montal, M., & Patapoutian, A. (2016). LRRC8 Proteins Form Volume-Regulated Anion Channels that Sense Ionic Strength. Cell, 164(3), 499–511. 10.1016/j.cell.2015.12.031

Szabadkai, G., Bianchi, K., Várnai, P., De Stefani, D., Wieckowski, M. R., Cavagna, D., Nagy, A. I., Balla, T., & Rizzuto, R. (2006). Chaperone-mediated coupling of endoplasmic reticulum and mitochondrial Ca2+ channels. The Journal of Cell Biology, 175(6), 901–911. 10.1083/jcb.200608073

Szymanowicz, O., Drużdż, A., Słowikowski, B., Pawlak, S., Potocka, E., Goutor, U., Konieczny, M., Ciastoń, M., Lewandowska, A., Jagodziński, P. P., Kozubski, W., & Dorszewska, J. (2024). A Review of the CACNA Gene Family: Its Role in Neurological Disorders. Diseases, 12(5), 90. 10.3390/diseases12050090

Voss, F. K., Ullrich, F., Münch, J., Lazarow, K., Lutter, D., Mah, N., Andrade-Navarro, M. A., von Kries, J. P., Stauber, T., & Jentsch, T. J. (2014). Identification of LRRC8 Heteromers as an Essential Component of the Volume-Regulated Anion Channel VRAC. Science, 344(6184), 634–638. 10.1126/science.1252826

Yanushkevich, S., Zieminska, A., Gonzalez, J., Añazco, F., Song, R., Arias-Cavieres, A., Granados, S. T., Zou, J., Rao, Y., & Concepcion, A. R. (2025). Recent advances in the structure, function and regulation of the volume-regulated anion channels and their role in immunity. The Journal of Physiology, 603(15), 4255–4291. 10.1113/JP285200

Zhou, X., Chen, Z., Xiao, L., Zhong, Y., Liu, Y., Wu, J., & Tao, H. (2022). Intracellular calcium homeostasis and its dysregulation underlying epileptic seizures. Seizure, 103, 126–136. 10.1016/j.seizure.2022.11.007

Zündorf, G., & Reiser, G. (2011). Calcium Dysregulation and Homeostasis of Neural Calcium in the Molecular Mechanisms of Neurodegenerative Diseases Provide Multiple Targets for Neuroprotection. Antioxidants & Redox Signaling, 14(7), 1275–1288. 10.1089/ars.2010.3359

